# The contribution of aquatic foods to human nutrient intake and adequacy in a Small Island Developing State

**DOI:** 10.1101/2024.10.02.616287

**Authors:** Jessica Zamborain-Mason, Jacob G. Eurich, Whitney R. Friedman, Jessica A. Gephart, Heather M. Kelahan, Katherine L. Seto, Neil L. Andrew, Michael K. Sharp, Aritita Tekaieti, Eretii Timeon, Christopher D. Golden

## Abstract

Many Small Island Developing States (SIDS) are experiencing a nutrition transition, wherein high prevalence of malnutrition co-occurs with growing rates of diet-related non-communicable diseases. Sustainably managed and accessible aquatic foods can serve as a rich and bioavailable source of nutrients, helping communities achieve healthy diets and meet key sustainable development goals (e.g., SDG 1 No Poverty, SDG 2 Zero Hunger, and SDG 14 Life Below Water). However, to properly harness aquatic food systems in nutrition interventions, we must first understand aquatic food’s role in nutrient intake and adequacy. Here, using a nationally representative survey from Kiribati, we quantify the contribution of aquatic foods to nutrient intake and adequacy, and examine the spatial variability in nutrient intake adequacies. We find aquatic foods are the main contributors of most nutrients we examined, providing > 80% of vitamin B_12_, retinol, and heme iron, and > 50% of niacin, vitamin A, protein, vitamin E, potassium, and total iron consumed. Consumption of aquatic foods contributes to meeting key nutrient adequacies (e.g., niacin) and provides complete adequacy for vitamin B_12_ and protein. However, despite high aquatic food consumption, we find high levels of nutrient inadequacies (11 of the 17 nutrients with dietary reference intakes). Overall, our study quantifies the nutritional importance of aquatic foods in an emblematic SIDS, emphasizing their vulnerability to declining aquatic resources. We also highlight the need for cross-scale context-specific targeted nutrition interventions, even when aquatic food consumption is high, to enable SIDS to meet key SDGs.

## INTRODUCTION

Tropical coastal communities around the globe are at the forefront of the triple burden of malnutrition, where undernutrition (e.g., stunting and wasting), overnutrition (e.g., obesity) and micronutrient deficiencies co-occur. Small Island Developing States (SIDS) are experiencing these dynamics [1,2]], with populations estimated to face high levels of micronutrient deficiencies [3,4], obesity (e.g., reaching >70% of the adult population in some locations, [5]), and stunting (e.g., as high as 51% [6]). Simultaneously, these populations are experiencing a nutrition transition towards highly processed, densely caloric, and sugar-sweetened foods that increase their risk of non-communicable diseases such as heart disease, cancer, chronic respiratory disease, and diabetes [7,8]. For example, a recent review covering 37 SIDS around the globe, revealed that one in six adults die prematurely from non-communicable diseases in over 80% of SIDS [6]. Increased malnutrition in such regions is driven by diets high in ultra-processed imported foods, and low in nutrient-dense foods [2,6,9]. Consumption of imported foods, which are often highly processed and contain high levels of harmful fats and sugars [10], has increased from 40% to over 60% in the last decade for both Pacific and Caribbean SIDS [6].

While aquatic foods are an important part of the diet for SIDS populations and billions of people globally, their role in nutrient intake and adequacy is not fully understood and is often undervalued [11,12]. Diverse aquatic foods are typically reduced to the oversimplified category of ‘fish’, and the diversity of nutrients provided is often ignored, with an overemphasis on protein or energy intake [13,14,15,16]. Aquatic foods are a rich, diverse, and bioavailable source of macro and micronutrients [17,11]and, compared to domesticated terrestrial animal-source foods, aquatic foods are a critical source of key nutrients such as omega-3 fatty acids, vitamin B_12_ and calcium [13]. In addition to providing key nutrients, aquatic foods can act as an alternative to less-healthy foods, consumption of which can lead to overnutrition-related health outcomes such as cardiovascular disease [18].

Realizing and maintaining the full nutritional and health benefits of aquatic foods is important for achieving key Sustainable Development Goals (SDGs) of SIDS [19,20]. For example, SDG 2 “*End hunger, achieve food security and improved nutrition and promote sustainable agriculture*” necessitates the availability of both adequate calories *and* nutritionally necessary macro and micronutrients [21]. Maintaining and/or increasing access to aquatic foods represents an opportunity for SIDS communities to consume adequate calories, and diverse micro and macronutrients, necessary to achieve the goals of SDG 2. Beyond direct nutritional benefits, aquatic foods represent an economic opportunity for many communities, delivering on SDG 1 “*No poverty*” and SDG 8 “*Decent work and economic growth*”. However, for aquatic foods to aid in meeting these SDGs, SDG 14 “*Conserve and sustainably use the oceans, seas and marine resources for sustainable development*” must also be met because without sustainable and resilient aquatic food resources, poverty, malnutrition, and unequal economic growth are likely to increase [21].

Strategies aimed at the sustainable and nutrition-sensitive management of aquatic food resources have been proposed to directly and indirectly help address the triple burden of malnutrition while helping advance other SDGs [22,23,16,19]. However, while it is typically assumed that aquatic food products are an important nutrient source for tropical coastal populations and SIDS, such contribution is seldom quantified [24,12]. Besides limited aquatic food group resolution in nutrient intake estimates, national apparent consumption of nutrients are often used as proxies for intake (e.g., [24]). These supply proxies linked to national exports and imports, are a measure of food availability, not actual intake, and, besides undervaluing small-scale and subsistence food production systems characteristic of many SIDS, they mask differences in aquatic food intake and reliance across groups (e.g., intra-national diversity). Knowing how aquatic foods, and different aquatic food groups, contribute to overall nutrient intake and the meeting of nutrient adequacy, and how this varies spatially, is critical to inform fisheries management plans and intervention programs aimed at improving nutrition and wellbeing of coastal tropical regions [12].

Here, using a nationally representative survey of dietary intake in Kiribati (including 2,146 households and 111 communities across 21 islands), we quantified the contribution of aquatic foods to overall nutrient intake and adequacy. Our study context is characteristic of many SIDS across the global tropics that are at the forefront of climate change and nutrition transitions [6]. Using household-specific nutritional intake of all food groups eaten, we quantify the contribution of aquatic foods and different aquatic food groups to dietary source nutrient intake. We then compare household intakes to household-specific nutritional requirements (Methods) to assess the nutrient intake adequacy of our study population, quantifying how aquatic foods contribute to meeting different nutritional inadequacies. Ultimately, we examine spatial variability in nutrient intake adequacy and aquatic food contributions at multiple scales, highlighting locations that either exceed or fall short from average nutritional conditions.

## RESULTS

### Quantifying the contribution of aquatic foods to household nutrient intake

Aquatic food products were not the major sources of total household food (in grams) or kilocalorie intake, contributing 19% of volume and 11% of dietary energy consumed by a household. However, they were the main source of vitamin B_12_ (cobalamine), both heme and total iron, retinol and overall vitamin A, vitamin B_3_ (niacin), protein, vitamin E, potassium, and calcium (i.e., 10 out of 21 nutrients we examined; Methods), and the second highest source of vitamin B_2_ (riboflavin), magnesium, zinc, vitamin B_1_ (thiamin), and sodium (i.e., totaling 15 of the 21 nutrients examined; Fig. 1a; Fig S1a). On average, aquatic foods provided ∼85% of household intake of vitamin B_12_, >75 % of heme iron, and retinol, and >50% of niacin and vitamin A (Fig. 1a). Aquatic foods also provided nearly half of household protein, >30% of vitamin E, riboflavin, magnesium, potassium, and >20 % of total iron, calcium, total fats, and zinc. Other foods such as bread and cereals were the main contributors of zinc, thiamin, magnesium, riboflavin, carbohydrates, and fiber, whereas fruits (of which coconut was the most common fruit consumed) were the main sources of non-heme iron, vitamin C, and total fats. Vegetables were the main source of household beta-carotene intake, and other food products not elsewhere classified (e.g., salt and soy sauce) were the main sources of sodium.

**Figure 1.**
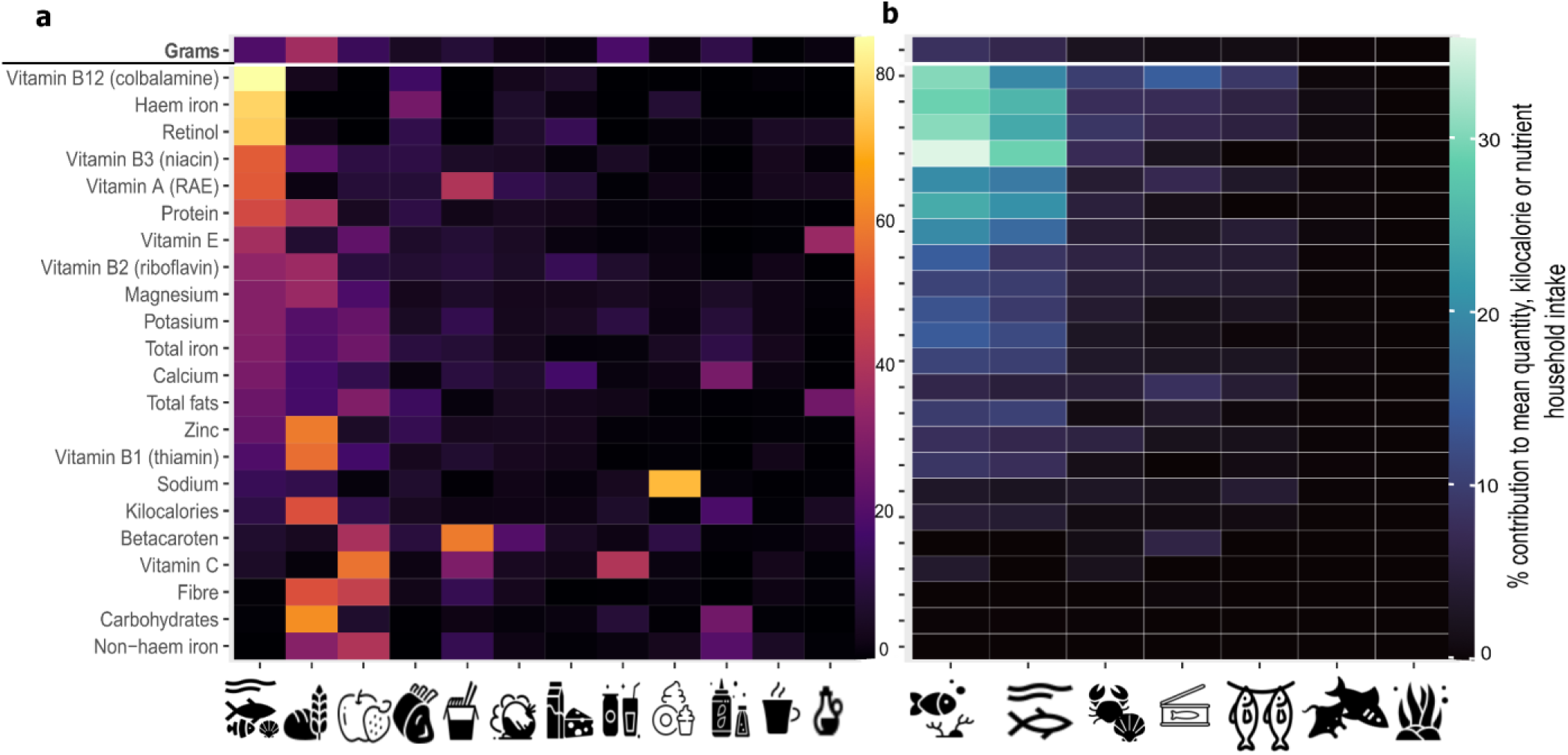
Contribution of aquatic foods to mean quantity (grams), kilocalorie and nutritional household intake. **(a)** Percent nutrient contribution of all different foods consumed. **(b)** Percent nutrient contribution of different aquatic food groups consumed. Food groupings on the x-axis are ordered based on their contribution to all nutrients combined. Besides grams, nutrients on the y-axis are ordered according to those that were most contributed by aquatic foods. Icons represent different food and aquatic food groupings, respectively: from left to right, **(a)** aquatic foods; bread and cereals; fruits; meat; restaurants, cafes and the like; vegetables; milk, cheese and eggs; mineral water, soft drinks, fruit and vegetable juices; sugar, jam, honey, chocolate and confectionary; Food products not elsewhere classified (e.g., salt and soy sauce); coffee, tea and cocoa; and oils and fats; **(b)** reef fish; pelagic and other fish; invertebrates; tinned fish; dried and salted fish; sharks and rays; and seaweed. See Fig. S1 for food group rankings instead of % contributions.

Disaggregating aquatic food contributions by aquatic food groups, we found that, in Kiribati, reef fish were the major contributor to nutrients obtained from aquatic foods, followed by pelagic and other fish (e.g., not those identified as reef fish, sharks or rays), invertebrates, tinned fish (e.g., tuna and mackerel), dried and salted fish, sharks and rays, and, lastly, seaweed. More specifically, based on mean household values, reef fish, besides being the major contributor to aquatic food kilocalories and grams of intake, was the main contributor to 15 of the 21 nutrients examined (excluding total fats, sodium, carbohydrates, calcium, betacarotene, and fiber; Fig S1). Reef fish accounted for 4% and 8% of the kilocalories and grams consumed by households, respectively, but contributed >30% of vitamin B_12_, and retinol,, ∼29% of heme iron, >20% to overall household vitamin A (RAE), protein, and niacin intakes, and >10% of vitamin E, riboflavin, magnesium, potassium, and total iron. Pelagic and other fish followed closely, contributing 4% and 7% to household intakes of kilocalories and grams, respectively, and contributing 29% of retinol intake, >20% of heme iron, and vitamin A, >15% of vitamin B_12_, niacin, and protein, and >10% of potassium, total iron, and total fats. Invertebrates, which only contributed 2% of the grams consumed by a household (and 0.9 % of kilocalories), provided >10% of household vitamin B_12_, and >5% of retinol, heme iron, vitamin A, vitamin E, and zinc. From all other aquatic food groups, tinned fish was the group that contributed most to household calcium intake (8%), and dried and salted fish contributed most to household sodium consumption (4%).

### Contribution of aquatic foods to household nutrient adequacy

To understand how nutrient intake from aquatic foods help meet nutrient intake adequacy, we compared nutrient intakes to household-specific recommended dietary intakes (e.g., recommended daily allowances), excluding four nutrients (i.e., heme iron, non-heme iron, retinol, and beta carotene) that do not have dietary intake thresholds defined (Methods). This revealed three key findings.

First, aquatic foods contribute to meeting key nutrient intake adequacies such as vitamin B_12_ and niacin (Fig. 2a). Consumption of aquatic foods alone provides more than triple the vitamin B_12_ requirements, enough protein (i.e., >100% requirements), >60% of niacin, >30% magnesium and sodium, and >20% of zinc, magnesium, potassium, riboflavin, and iron (Fig. 2b). Without aquatic foods (assuming they are removed from the diet and not replaced by other foods), our study population would only meet nutrient intake requirements of carbohydrates, sodium, protein, and vitamin C.

**Figure 2.**
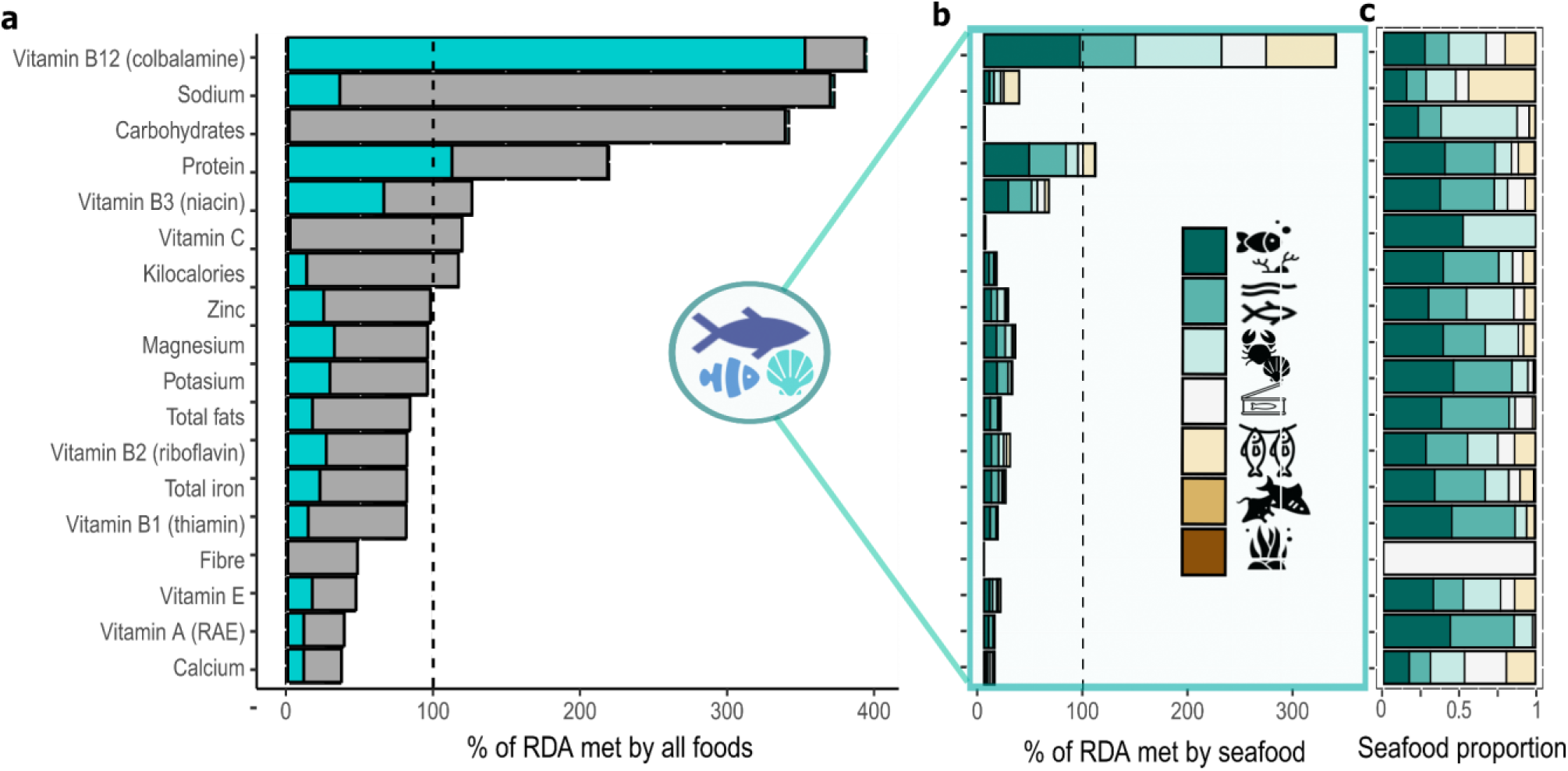
Contribution of aquatic foods to mean kilocalorie and nutritional household adequacy (% of household recommended daily allowance met). **(a)** Overall from all food groups eaten, separating aquatic foods’ contribution (turquoise) from all other foods (dark grey), **(b)** only from aquatic foods, and **(c)** proportion of aquatic foods’ adequacy separated by aquatic food groups. Colors in **(b)** and **(c)** represent different aquatic food groups shown in the legend as icons, from top to bottom: reef fish; pelagic and other fish; invertebrates; tinned fish; dried and salted fish; sharks and rays; and seaweed. Nutrients on the y axes are ordered by mean nutritional household adequacy from all food groups eaten.

Second, even under our study context, which, on average, has high levels of aquatic food consumption (Fig. S2) and meets kilocalorie recommendations, there were still high levels of inadequate nutrient intake (Fig. 2a). More specifically, from the 17 nutrients that had dietary reference intakes, on average, our study population only met six (i.e., vitamin B_12_, sodium, carbohydrates, protein, niacin, and vitamin C), with intake of three (i.e., vitamin B_12_, sodium, and carbohydrates) being triple the requirement. On average, the population did not meet nutrient intake adequacy of total fats, vitamins A, E, B_2_ (riboflavin), B_1_ (thiamin), minerals such as calcium, iron, zinc, magnesium, and potassium, and dietary fiber.

Third, examining the contribution of specific aquatic food groups, we found that while reef fish contributed the most to aquatic food nutrient intake adequacy across nutrients, other groups such as invertebrates or pelagic fish disproportionately contributed to specific nutrients such as zinc or total fat adequacies, respectively (Fig. 2c). More specifically, consumption of reef fish alone provided >95% of vitamin B_12_ requirements, >45 % of protein, >20 % of niacin, and >10% of magnesium and potassium requirements. Reef fish contributed 7% to zinc adequacy and 6% to total fat adequacy, while invertebrates and pelagic fish contributed slightly more (i.e., 8%) to both zinc and total fat adequacies.

### Variability in aquatic food contributions to nutrient intake adequacy

Overall, we found high spatial variability across households, villages, and islands in Kiribati of the prevalence of meeting nutrient intake adequacies and aquatic foods’ particular contribution to nutrient intake and adequacy (Fig. S3-S4). We also found inter- and intra-island spatial variability in the types of aquatic foods consumed, with some households, for example, obtaining 100% of their aquatic food nutrients from single groups such as invertebrates (Fig. S3). Thus, accounting for the nested structure of our data (i.e., households within villages within islands; see Methods), we (i) quantified spatial variability in nutrient intake adequacy for our study population, as well as in the contributions of aquatic foods to meeting such adequacy, and (ii) examined whether there were locations meeting higher levels of nutrient intake adequacy. We found that the adequacy of micronutrients such as vitamin C and vitamin A varied more than the adequacy of macronutrients such as protein and carbohydrates (Fig. S5). We also found that spatial variability in nutrient intake adequacy and aquatic food contributions was always greatest among households within villages, our lowest unit of analysis, in comparison to among islands or among villages within islands (Fig. S5).

However, perhaps most importantly, we identified locations at all levels (i.e., households, villages, and islands) within our study region that had significantly higher levels of nutrient intake adequacy. >85 % of households consumed adequate amounts of carbohydrates, protein, vitamin B_12_ and sodium, >50% households met kilocalorie and niacin requirements, > 30% consumed sufficient vitamin C, zinc, potassium and magnesium, >20 % met fat, riboflavin, iron and thiamin, and <10 % of households met requirements for vitamin A, fibre, vitamin E and calcium. Only 14 households (0.7% of households examined) met all nutritional intake adequacies (i.e., kilocalories and 17 nutrients), however, these were spread across 12 different villages and eight islands. While no island or village met all nutrient intake adequacies, some locations deviated positively and negatively from average nutrient intake adequacy conditions (Fig. S6-S7). Some islands (e.g., Nonouti, Butaritari, and Beru), for instance, consistently had lower levels of adequacy across all nutrients and kilocalories (Fig. S6). Others, however, deviated from country-level adequacy status positively for some nutrients and negatively for others. South Tarawa, for example, which is the country capital, had higher levels of sodium, carbohydrates, fats, niacin, zinc, thiamin, riboflavin, iron, fiber, and calcium adequacy, but lower levels of vitamin B_12_, potassium, and vitamin C. Kiritimati, on the other hand, deviated from average national-level conditions positively for kilocalories, sodium, protein, vitamin B_12_, niacin, zinc, thiamin, riboflavin, vitamin E, vitamin A, and calcium, and negatively for fiber and vitamin C (Fig. S6). Within islands, there were also villages that deviated from island level averages (Fig. S7). For example, within Kiritimati, London had higher sodium intake and lower carbohydrates, vitamin B_12_, protein, zinc, vitamin C, and vitamin A adequacy levels than neighbouring villages (i.e., Poland, Banana, and Tabwakea).

Aquatic food contributions to nutrient intake adequacy also varied spatially but spatial patterns were much more consistent across nutrients (Fig. S5-S8). For example, islands such as Teeraina, Tabuaeran, North Tabiteuea, Marakei, Maiana, Kiritimati, Abemana, and Abaiang, obtained higher levels of nutrient intake adequacy from aquatic foods whereas islands such as South Tarawa, Nonouti, Nikunau, Butaritari, and Beru received significantly lower levels (Fig. S8). Similarly, specific villages within islands (e.g., Poland and Banana within Kiritimati) had higher contributions from aquatic foods to nutrient intake adequacy than their neighboring villages (e.g., London; Fig. S9). Looking at spatial variability in aquatic food group contributions, our study shows that islands such as Teeraina, Tabuaeran, Marakei, Kuria, Aranuka, Abemana, and Abaiang obtain higher levels of nutrient intake adequacy from reef fish (Fig. S10), whereas islands such as Onotoa, Kiritimati and also Marakei and Kuria deviate from average national conditions because they obtain more nutrients from pelagic fish (Fig. S11). Teeraina, Marakei, and Abemana stand out as consuming more nutrients from invertebrates (Fig. S12), and Beru, Kiritimati, and Teerania stand out, in comparison to other islands, as obtaining more aquatic food nutrients from tinned fish (Fig. S13).

## DISCUSSION

Our study shows that aquatic food intake is a critical source of nutrients for SIDS such as the Republic of Kiribati. On average, aquatic foods provide these populations >80% of vitamin B_12_, retinol, and heme iron, and > 50% of niacin, vitamin A, protein, vitamin E, potassium, and total iron consumed. This intake contributes significantly to people’s nutritional needs, preventing micronutrient deficiencies and non-communicable diseases. However, our findings also reveal that even in populations with high aquatic food consumption–a typical characteristic of our study population and many SIDS–there can still be significant instances of both overconsumption (e.g., carbohydrates and sodium) and underconsumption (e.g., iron, vitamin A, and calcium) of specific nutrients. Together, these results emphasize that (i) SIDS populations are highly vulnerable to declining aquatic resources or reduced access to aquatic foods, and (ii) even in the presence of sufficient aquatic foods, nutrition-sensitive approaches and targeted nutrition interventions are needed to enable SIDS to tackle malnutrition and NCDs.

Small-scale fisheries supply critical nutrients for coastal populations around the globe [24]. Historically, the contributions of aquatic foods to diets have primarily been restricted to protein (e.g.[25]), however we find that the importance of protein in seafood pales in comparison to the importance of key micronutrients such as vitamin B_12_, retinol, and heme iron, contributing towards meeting intake recommendations of SIDS populations. In fact, even in systems where aquatic foods are the main animal-source food, populations can already meet their protein requirements without aquatic food consumption, (e.g., Fig., 2). Typically, when households are meeting their caloric needs, they tend to be obtaining adequate protein (Fig. S14). It is these other nutrients that are particularly important in the absence of, or reduced access to, other animal-source foods, as is characteristic of many SIDS [20]. Vitamin B_12_, for example, is critical for cognitive function and intestinal immune regulation [26], niacin plays a key role in neural development and overall survival [27], and heme iron, which is better absorbed by the body in comparison to non-heme iron, is critical for transporting oxygen to cells [26] with iron deficiency anemia linked to higher levels of fetal death and maternal morbidity [28].

Declines in aquatic food intake in SIDS, due to reduced availability [29] or reduced access [30,19], can put populations at higher risk of nutritional deficiencies, and further increase their vulnerabilities [31]. This is particularly relevant for aquatic food groups that contribute significantly to people’s nutrient intake adequacy, such as reef and pelagic fish (Fig. 2), and that are changing due to environmental and anthropogenic stressors. Reefs, for example, are functionally changing or declining worldwide due to factors such as thermal stress [32,33]and overfishing [34], whereas pelagic resources such as tuna are projected to redistribute under climate change and become less accessible for many SIDS [35]. Thus, increasing the sustainability and climate-resilience of local aquatic resources (e.g., through effective and equitable climate-resilient fisheries management; [36]), as well as reinforcing access to locally available aquatic food products [19] should be a priority for SIDS aiming to improve malnutrition. In the absence of aquatic foods, populations could be immersed in low quality under-nourishing or over-nourishing diets (e.g., high in sodium, carbohydrates; Fig.2; [20,22]), further accelerating the nutrition transition and leading to poor health outcomes [37].

However, aquatic foods may not be a panacea to address all nutritional deficiencies. We show high levels of inadequacy for some nutrients (e.g., calcium, fibre, iron, vitamin A, and vitamin E; Fig. 2), even under the high levels of aquatic food consumption that are characteristic of many SIDS [12,20,38]. Calcium deficiency is linked to low bone mass, osteoporosis, and preterm birth [39]. Fibre deficiency, on the other hand, increases the risk of cardiovascular disease and colon cancer [39], whereas deficiencies of vitamin A and E can cause increased susceptibility to infections [26]. This highlights the need to consider the food system as a whole, and the benefit that public health interventions aimed at increasing food diversity [12], access to other nutrient-dense foods (e.g., nuts, fruits, and vegetables), and incorporating effective nutrient fortification and supplementation in diets (e.g., [40]) can have on the health and wellbeing of people living in SIDS. Improved nutrient intake adequacies can be achieved, for instance, by promoting healthier imports [41] or the consumption of products rich in nutrients of public health concern. Some nutrient intake adequacies may also be achieved by increasing the consumption of locally available underutilized foods such as seaweed. In comparison to animal-source aquatic foods, which dominate current consumption patterns (Fig. 2), seaweed has high concentrations of fiber, magnesium, iron, and vitamin E [17], nutrients that are currently consumed in insufficient quantities by these populations. Exploring the potential of locally available solutions like incorporating seaweed in people’s diets may be critical for SIDS that have limited land, freshwater resources, and agricultural production [2].

Ultimately, our spatial variability results highlight that, within a country, there can be cross-scale positive examples (e.g., households, villages and islands) of meeting higher levels of nutrient intake adequacies, but also locations with higher levels of inadequacy that could be prioritized for nutrition interventions. As expected, given the quantities of nutrients in foods, we found that spatial variability was greater for micronutrient intake rather than kilocalorie or macronutrient intake (e.g., Fig S5; [42]). However, we also found spatial variability was greatest at our lower scale of analyses (i.e., households within villages). While this variation could be partially driven by the natural week-to-week variation of people’s diets [42], it also suggests that besides global and regional processes affecting access to food resources and nutritional outcomes (e.g., globalization, development or environmental degradation;[1,22], household and individual socio-behavioural characteristics (e.g., income, social status, or food choices) may be key drivers shaping nutrient intake adequacy [19,43]. Future research could build upon our work to discern the diverse environmental and socio-economic drivers of meeting nutrient intake adequacies across scales, and the drivers of aquatic food contributions, and thus help find locally applicable management and policy opportunities to tackle malnutrition. For example, village and island level differences may showcase where infrastructure, education, trade or geographical access play an important role in dietary behaviors, or macro-level dynamics inhibiting intake of substantial aquatic food-based nutrients in some locations (e.g. Nonouti, Butaritari, Beru) Furthermore, work that incorporates temporal variability (e.g., multiple food recalls in different seasons; [42]) and intrahousehold variability (e.g., dietary intake methods that allow for gender and age differences;[4,44,45,46]) will also be critical to determine temporal and demographic variabilities in nutritional outcomes.

Many SIDS around the globe are at the forefront of nutrient-transitions. While addressing issues within the aquatic food production system is recognized as a key component of nutrition and development interventions across SIDS, a deeper understanding of aquatic foods’ role in nutrient intake and adequacy, and their variability, is essential for enabling these communities to meet sustainable development goals, such as SDG 1 (No Poverty), SDG 2 (Zero Hunger), and SDG 14 (Life Below Water). Here we show that maintaining access to local aquatic foods is critical to avoid populations further immersing in undernourishing and overnourishing diets, the characteristics of a nutrition transition. However, the vulnerability of these populations to declining aquatic resources depends on the effective implementation of multidisciplinary strategies aimed to enhance food system sustainability and resilience. Effective fisheries management that enhances the sustainability, resilience and accessibility of local aquatic foods will help people meet key nutrient intake adequacies. However, to tackle the triple burden of malnutrition, such a strategy needs to be coupled with local, national and international interventions that ensure people also have access to other nutrient-dense foods, that combined with aquatic foods create healthy and nutrient-rich dietary patterns. By quantifying the nutritional intake and adequacy across multiple scales, and how aquatic foods contribute towards that, we show the importance of cross-scale context-specific interventions. Future work that unlocks the multiscale drivers of nutritional adequacy in such regions will be critical for communities and nations to effectively navigate the triple burden of malnutrition and enhance the well-being of their populations.

## METHODS

### Study system

We quantified the nutritional role of aquatic foods for SIDS using the Republic of Kiribati, an emblematic SIDS [6], as a case study. The Republic of Kiribati is an independent nation in the Central Pacific that has 33 low-lying islands distributed in three major island groups: the Gilbert Islands, the Phoenix Islands, and the Line Islands. Combined, Kiribati has a total land area of 811 km^2^ and ∼3.55 million km^2^ of ocean area [47], representing the largest ocean-to-land ratio in the world. Due to the available resources and geographic isolation of the country, seafood consumption in Kiribati is one of the highest in the world, with high dependence on reef-based resources [43]. Moreover, shifts in economic conditions, environmental factors, and climate change are likely to jeopardize food availability, income generation, and the health of local ecosystems, which could, in turn, impact the nutritional well-being and disease prevalence among the I-Kiribati population [22].

We conducted our research in 21 of the 23 inhabited islands of Kiribati across the Gilbert and Line Islands groups structured around a 2019-2020 nationally representative, observational cross-sectional social study called the Household Income and Expenditure Survey (HIES; see [48]). These geographically and socially diverse surveys were implemented by the Pacific Community in partnership with the Kiribati national government. Primary respondents self-identified as the person in the household with the most knowledge about household food consumption. Full information and questionnaire for the May 2019 to March 2020 Kiribati HIES used here can be obtained from the Pacific Community (SPC [49]) (https://pacificdata.org/data/dataset/spc_kir_2019_hies_v01_m_v01_a_puf).

### Estimating the contribution of aquatic foods to household nutritional intake

To estimate the contribution of aquatic foods to household nutritional intakes we used household weekly food recalls collected through the HIES. Weekly food recalls were collected in a total of 2,146 households from 111 different villages across the 21 surveyed islands. Household members were asked if any member in the household had consumed specific foods within broad categories and the quantity consumed. Foods were divided into broad categories (e.g., grain and cereals; meats; aquatic foods; dairy and oils; fruits; vegetables and crops; beverages; snacks, candy and confectionery, spices and condiments; or prepared meals) and within broad categories, respondents could specify food consumed from a preset list and other foods not originally listed. Food consumed by the household was included, whether it was sourced from cash transactions, own-account production, gifting, or through exchange [48] To estimate the nutritional contribution of different aquatic food groups, within aquatic foods, we grouped food groups into seven categories: reef fish (e.g., snappers, groupers, emperors, parrotfish and surgeonfish), pelagic and other fish (e.g., tuna, deep water snapper and flying fish), sharks and rays, invertebrates (e.g., coconut and mud crabs, cockles, clams, sea snails, sea worms, and lobsters), seaweed (e.g., *Kappaphycus* spp.), dried/salted fish, and tinned fish (e.g., tuna and mackerel).

The Pacific Nutrient Database [50]was used to assign nutrient composition estimates to food recall data. PNDB was specifically developed to address the need for a standardized method for linking data between the HIES and nutrient composition data for foods consumed in the region [12]. The PNDB matched food items from the HIES with their respective macro- and micro-nutrient composition, presenting values per 100-gram edible portions for total energy, all macronutrients (carbohydrates, protein and total fats), fiber, and 17 micronutrients: retinol, beta carotenes, vitamins A (Retinol Activity Equivalents; RAE), B_1_ (thiamin), B_2_ (riboflavin), B_3_ (niacin), B_12_ (cobalamine), C and E, and sodium, magnesium, potassium, calcium, zinc, heme iron, non-heme iron, and total iron. Note that retinol differs from vitamin A (RAE), because the latter includes plant-based vitamin A sources such as beta carotenes. While not all nutrients important in aquatic foods are represented in the PNDB (e.g., long-chain omega-3 fatty acids; [17]), and edible portions could underestimate consumption of some nutrients (e.g., calcium if for example fish are eaten whole; [46]), we used this database because it is currently the most relevant for foods consumed in the region.

Household daily nutrient intake (for all foods combined, all aquatic foods, and individual aquatic food categories) was estimated by dividing weekly values by seven (i.e., seven days in a week). Contributions of aquatic foods, and different aquatic food groups, were contrasted to contributions from all food groups as percentages (e.g., contribution of aquatic foods to kilocalorie intake is equal to the ratio of kilocalories obtained from aquatic foods divided by the total kilocalories consumed, all multiplied by 100).

### Estimating the contribution of aquatic foods to nutrient intake adequacy

Household-specific nutritional requirements were estimated based on household-specific demographics (i.e., age, sex and, when relevant, pregnancy status) and demographic-specific nutrient recommended dietary allowances or adequate intakes [51]. As total fats recommended dietary allowances are not determined beyond infant categories, we calculated recommended total fats for the remaining demographic categories as 25% of recommended kilocalorie intakes [52], converting fat consumption to energy intake in calories (i.e., 1g of fat=9 Kcal; [53]). Due to the absence of breastfeeding data, the estimated household requirements are conservative. Specific nutrients available within the PNCD (e.g., heme iron, non-heme iron, retinol, and beta carotene) were excluded from this section due to a lack of recommended dietary allowances or adequate intake values. We compared household-specific apparent nutrient intake estimates to these as ratios (i.e., household apparent consumption divided by household requirements), whereby the value of one indicates the household met household-specific dietary requirements.

### Quantifying spatial variability in nutrient intake adequacy

To characterize spatial variability in nutrient intake adequacies, while controlling for the nested structure in our data, we separately modeled each outcome (i.e., overall nutrient adequacy, nutrient intake adequacy from aquatic foods, and nutrient intake adequacy from different aquatic food groups for each nutrient) using Bayesian multilevel models. We used lognormal family distributions for outcomes that did not have zeroes, and hurdle lognormal distributions for outcomes containing zeroes. Note that some nutrients were not modelled for specific aquatic food groups due to not having variability (i.e., all values being zero; e.g., vitamin C or fibre). All analyses were performed using the Hamiltonian Monte Carlo algorithm implemented in RStan [54] through the brms package in R [55]. Four chains were run for each scenario, leaving 4000 samples in the posterior distribution of each parameter. Convergence was monitored by running four chains from different starting points, examining posterior chains and distribution for stability, and checking that the potential scale reduction factor (also termed R_hat) was close to 1 (below 1.01; [56]). All models fit the data well (Fig. S15).

## ACKNOWLEDGEMENTS

This study was supported by National Science Foundation Award #1826668 CNH-L Interactive Dynamics of Reef Fisheries and Human Health (CDG, KLS, JAG, JGE, JZM and WRF). MS and NA were funded by the Australian Government through Australian Center for International Agricultural Research (FIS/2018/155). We are grateful to the i-Kiribati enumerators and teams that ran the HIES, our Kiribati MHMS colleagues Tebwebweiti Tikanibwebw, Tebano Bwabwa, Nantebwebwe Toabo, Bwaturia Temaua, Baurina Kaburoro, Tirite Irooti, Regina Flood, and Kiaman Raurenti for their collaboration with the HIES and food recalls, Douglas McCauley for research planning, and Kelvin Gorospe for data processing and structuring. Icons were obtained from icons8.com

## AUTHOR CONTRIBUTIONS

JZM conceived and developed the study with support from CG, KS, JG, JE and WF. MS helped with data acquisition, development and interpretation. JZM implemented the analyses. JZM wrote the first draft of the manuscript with help from HK. All authors contributed substantially.

## DATA AVAILABILITY STATEMENT

The data in this study were accessed by a use agreement with the primary custodians of the data, the Pacific Community (SPC) and the Government of Kiribati NSO. Requests for full data access can be made through SPC and Kiribati NSO offices. Additionally, data cannot be fully anonymized, as some villages have populations too small for true anonymization.

## SUPPLEMENTARY INFORMATION

**Figure S1.**
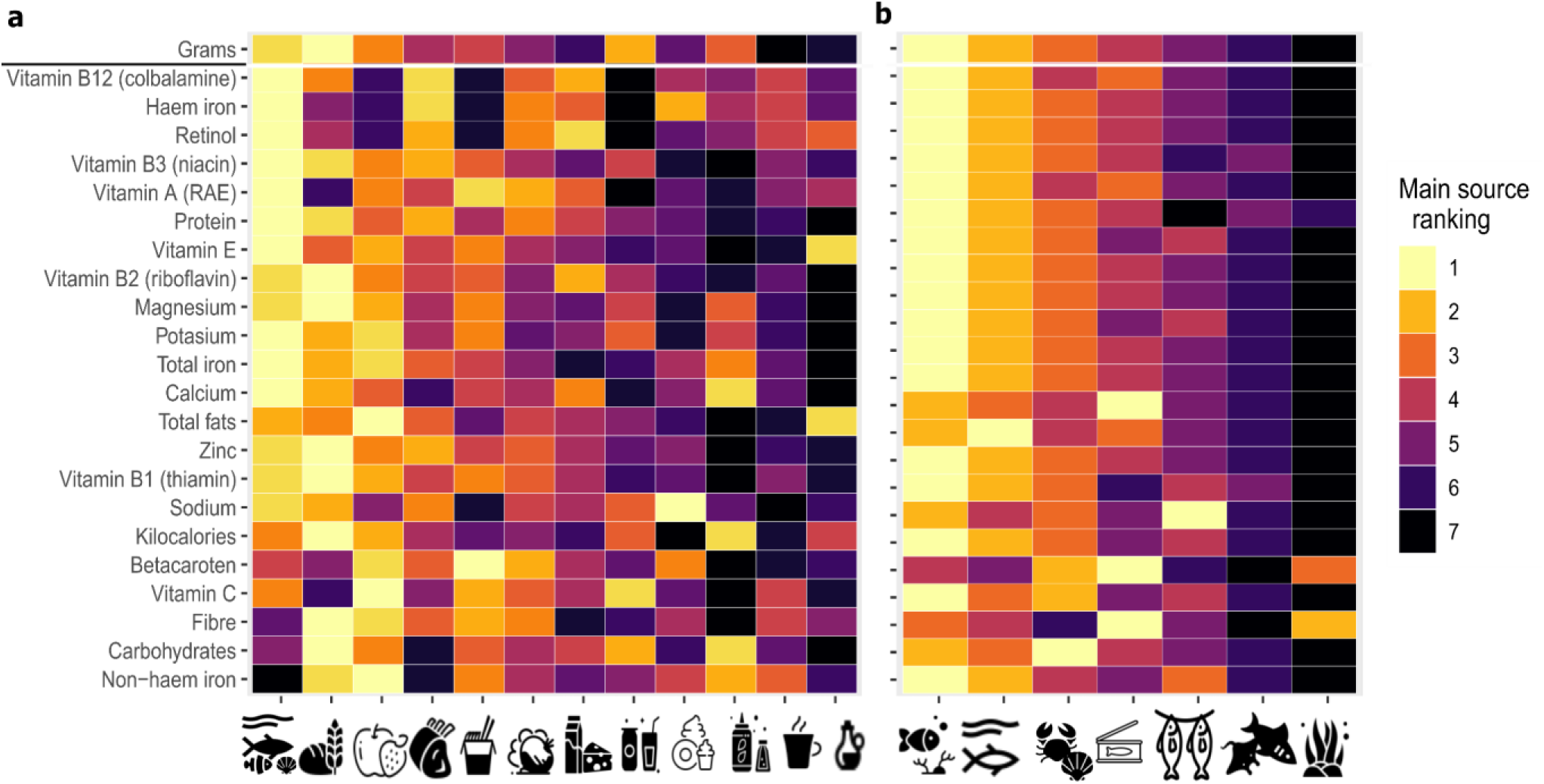
Ranking of aquatic foods based on their contribution to mean quantity (grams), kilocalorie and nutritional household intake. (a) Ranking based on the mean percent nutrient contribution of different foods consumed. (b) Ranking based on mean percent nutrient contribution of different aquatic food groups consumed. Food groupings on the x-axis are ordered based on their contribution to all nutrients combined. Besides grams, nutrients on the y axes are ordered according to those that were most contributed by aquatic foods. Icons represent different food and aquatic food groupings, respectively: from left to right, **(a)** aquatic foods; bread and cereals; fruits; meat; restaurants, cafes and the like; vegetables; milk, cheese and eggs; Mineral water, soft drinks, fruit and vegetable juices; sugar, jam, honey, chocolate and confectionary; Food products n.e.c (e.g., salt and soy sauce); Coffee, tea and cocoa; and oils and fats; **(b)** Reef fish; pelagic and other fish; invertebrates; tinned fish; dried and salted fish; sharks and rays; and seaweed. See Fig. S1 for rankings instead of % contributions.

**Figure S2.**
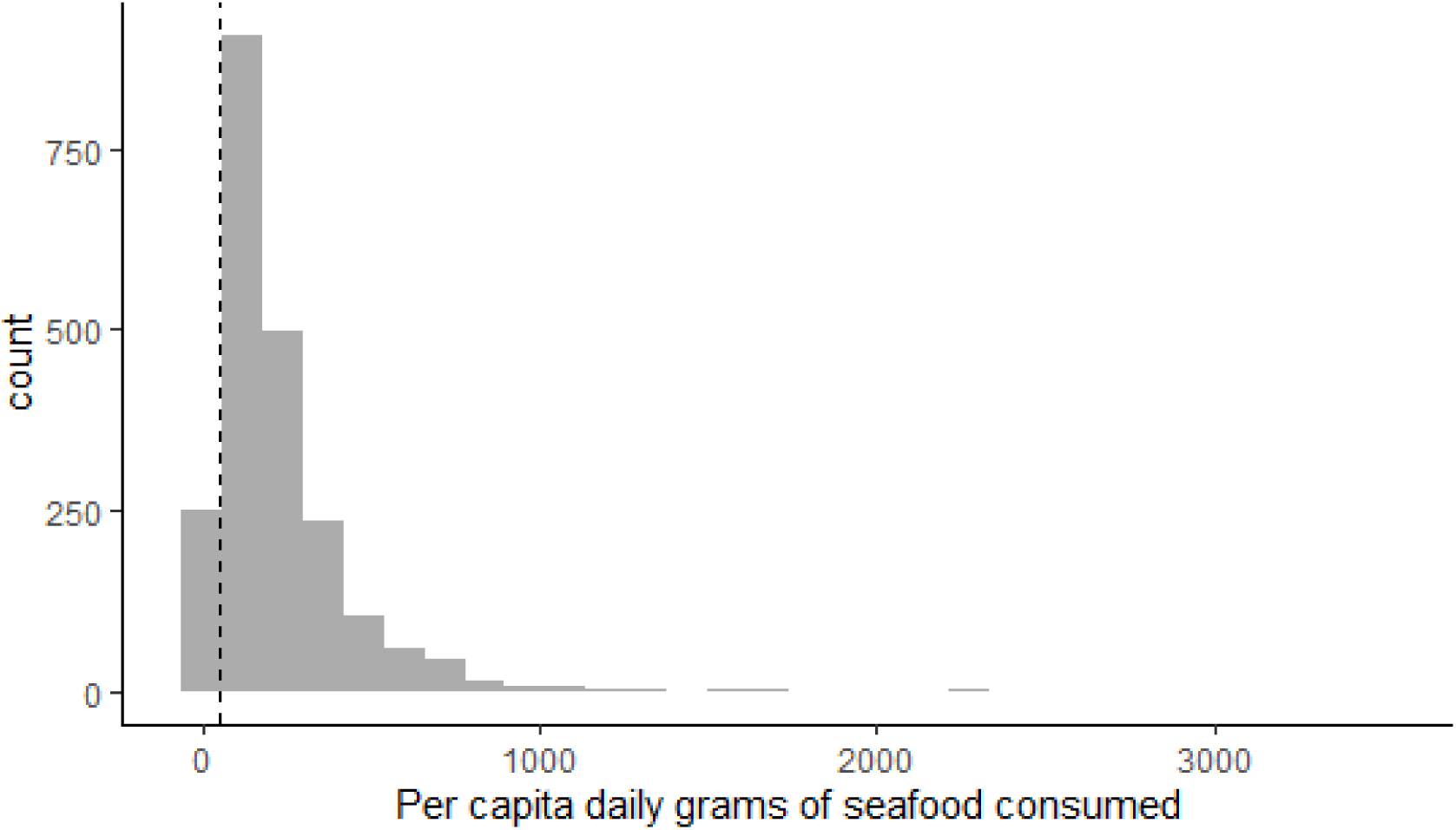
Estimated grams of aquatic foods consumed per day per capita based on household intake and household size. The distribution is based on different households, assuming an even spread of aquatic food consumption among household members. As a reference, the dashed vertical line represents 48.6 g-the recommended upper limit for pregnant women to minimize exposure to methylmercury based on 340 weekly grams [57].

**Figure S3.**
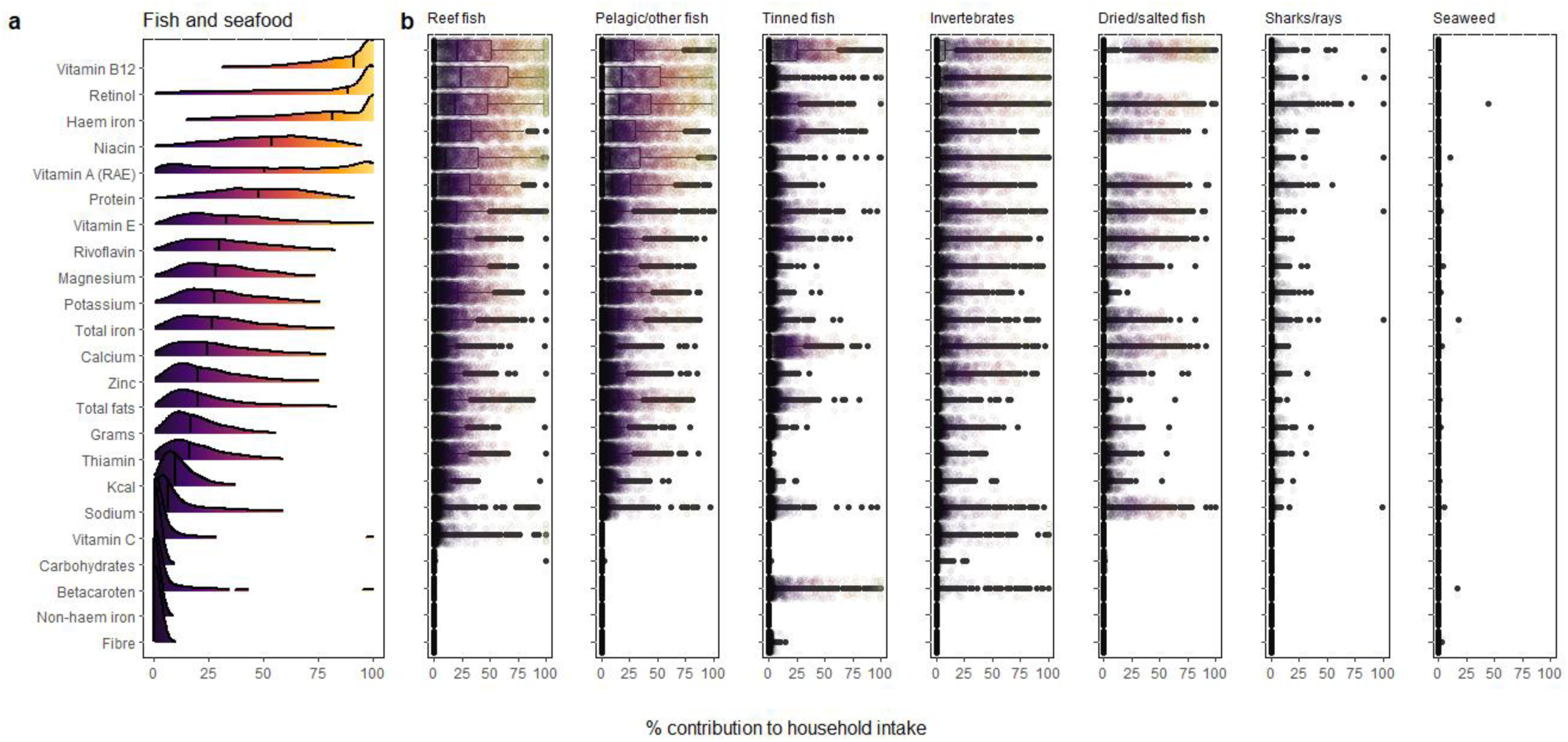
Observed variability in aquatic food contribution (a), and different aquatic food groups (b), to household nutritional intakes. Individual points in **(b)** represent households Vertical lines represent median values.

**Figure S4.**
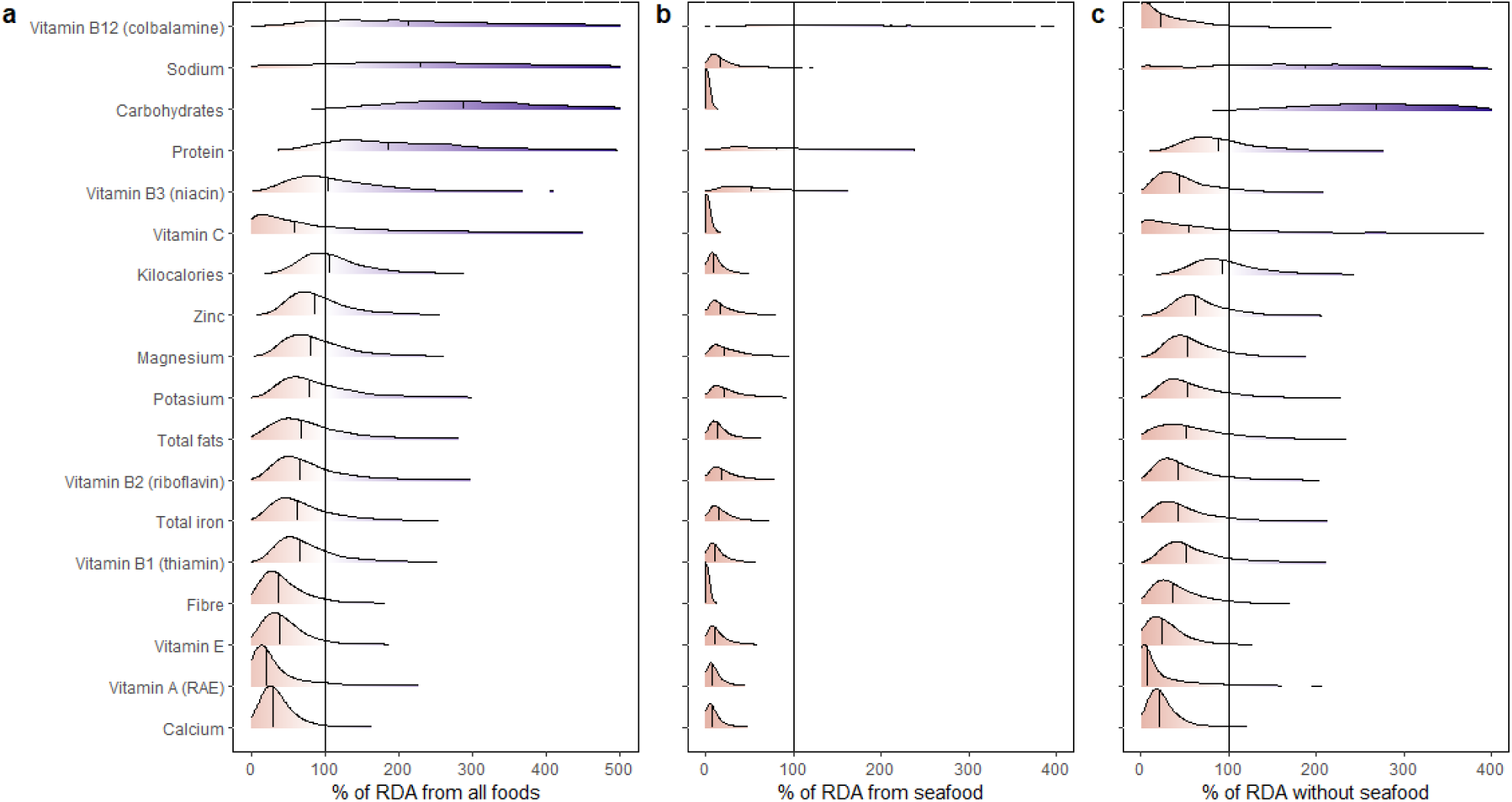
Observed variability in household nutrient intake adequacy (a), showing contributions only from aquatic foods (b) and without aquatic foods (c). Nutrients are ordered based on mean household values. Individual vertical lines on top of each density represent median values. Solid vertical lines represent 100, meaning nutritional intake for that household matched household-specific nutritional requirements.

**Figure S5.**
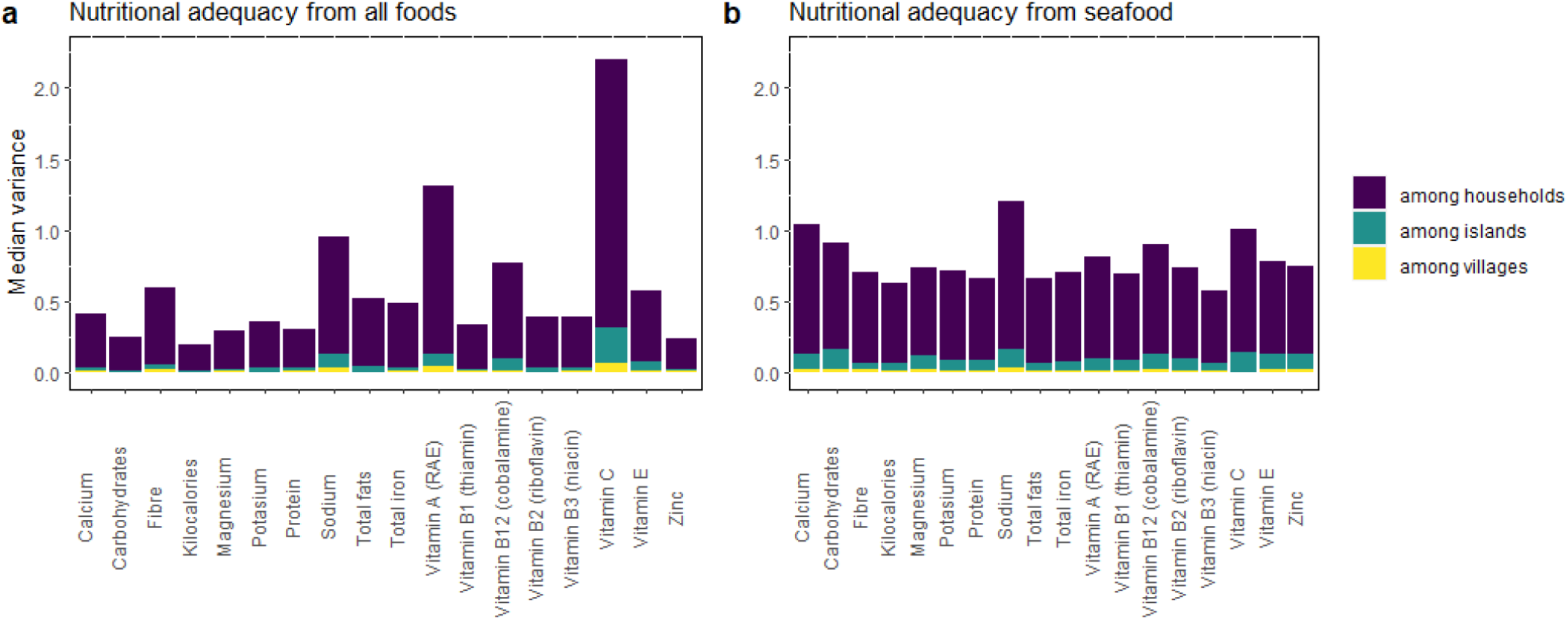
Variance decomposition of nutrient intake adequacy from all foods and only from aquatic foods separated by the nested structure of our data: among islands, among villages within islands, and among households within villages). Values are the median estimated variance from our bayesian hierarchical models.

**Figure S6.**
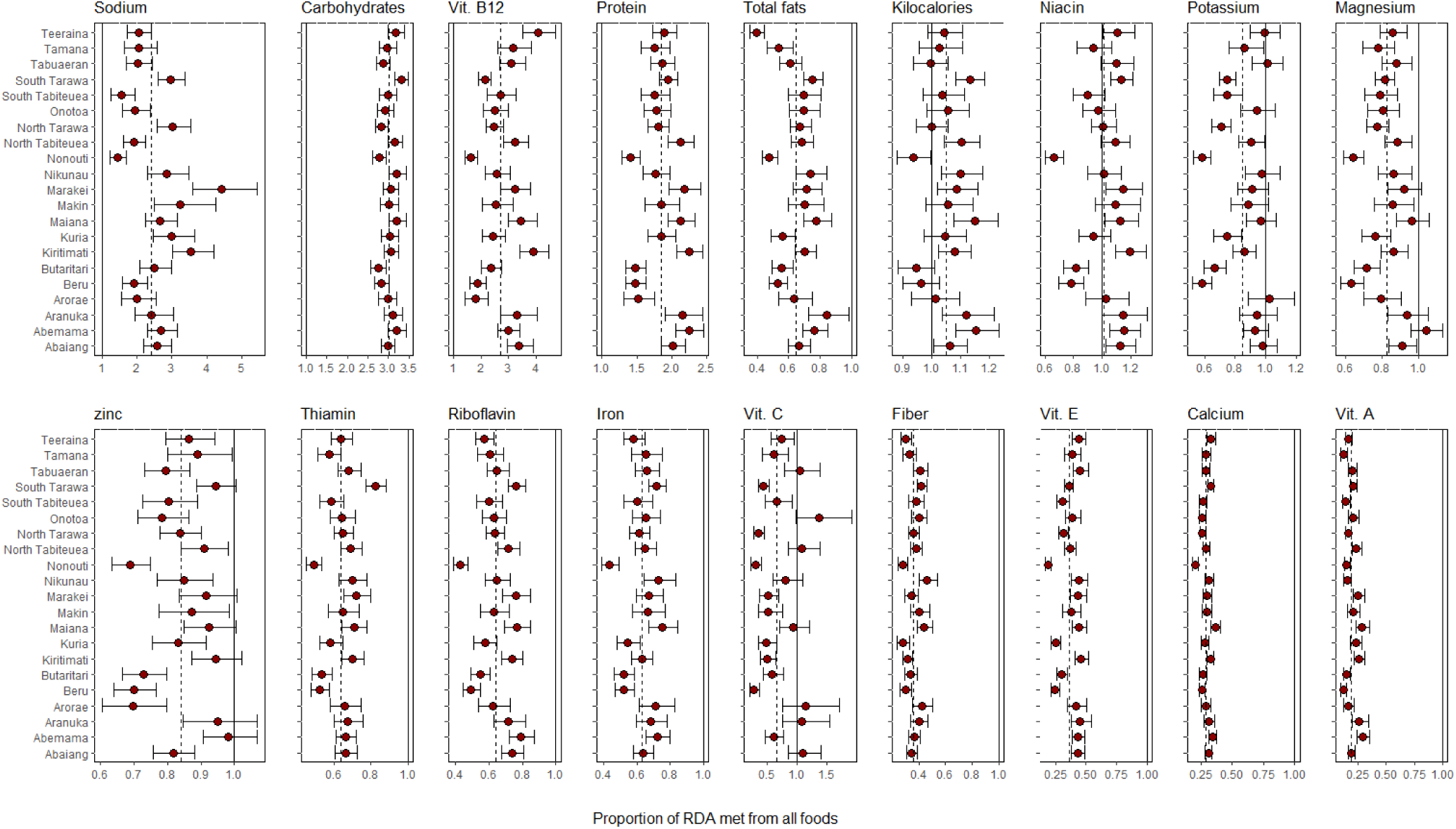
Island specific variability in nutrient intake adequacy(proportion of nutrient-specific recommended daily allowance met from all foods). Points are median estimated nutrient intake adequacy and intervals are 90% uncertainty intervals. Vertical lines indicate meeting adequacy (solid line=1) and average conditions for the entire population accounting for the nested structure in the data (dashed line; i.e., country-level intercepts).

**Figure S7.**
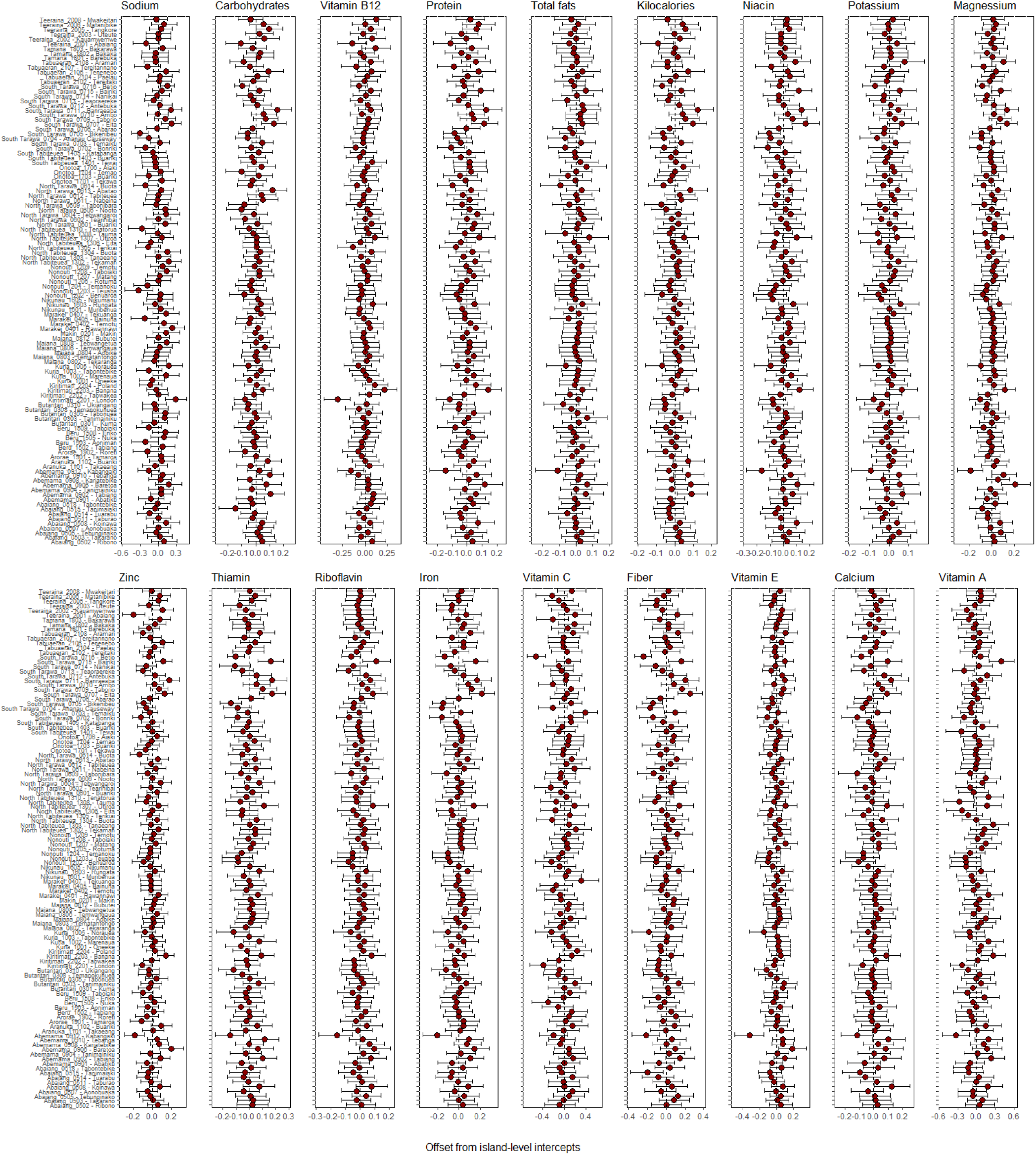
Community specific variability in nutrient intake adequacy as an offset from island-level estimated intercepts. Points are median estimates and intervals are 90% uncertainty intervals. Vertical dashed lines indicate island-level intercepts.

**Figure S8.**
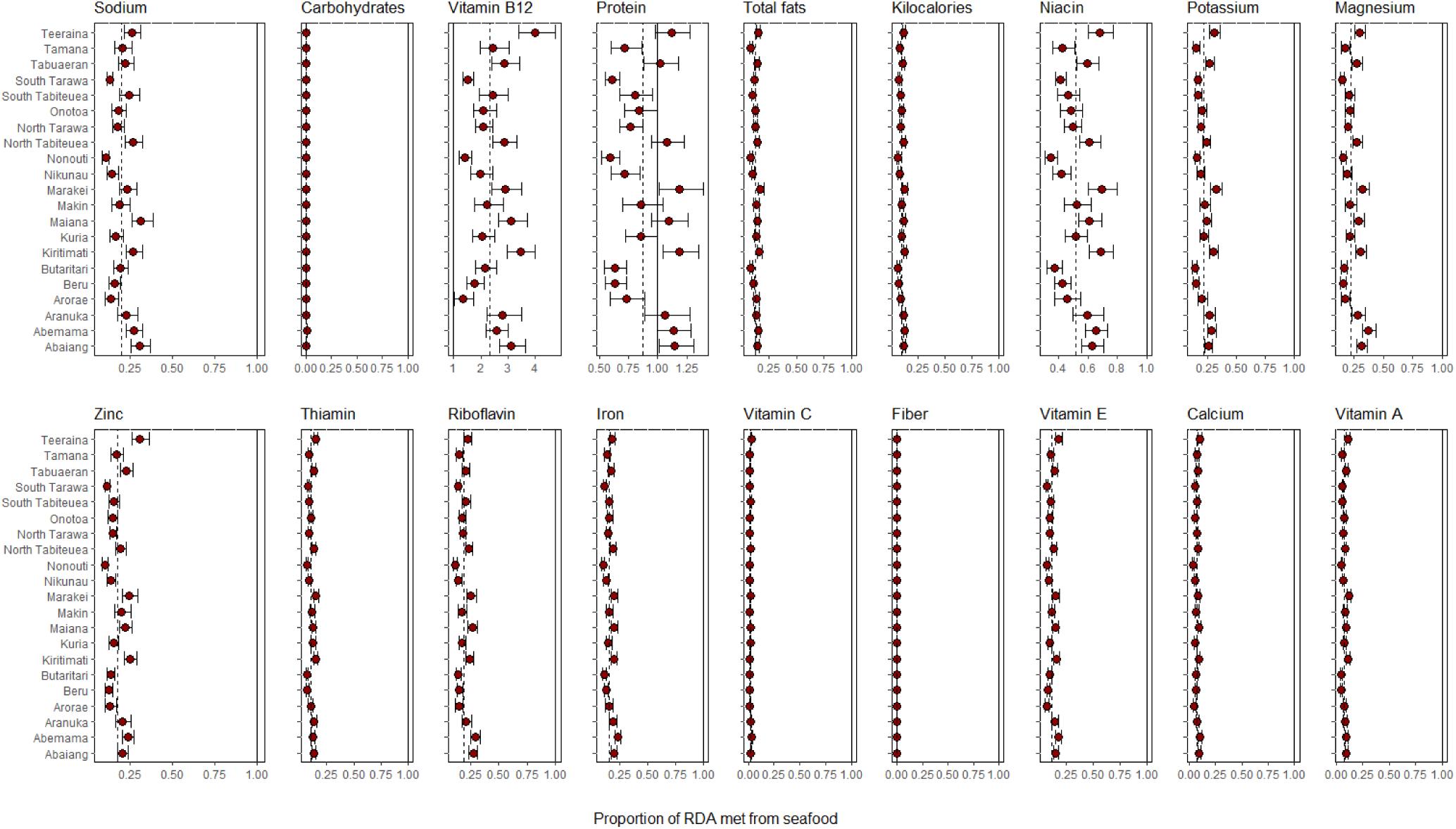
Island specific variability in nutrient intake adequacy from aquatic foods (proportion of nutrient-specific recommended daily allowance met from only aquatic foods). Points are median estimated nutrient intake adequacy and intervals are 90% uncertainty intervals. Vertical lines indicate meeting adequacy (solid line=1) and average conditions for the entire population accounting for the nested structure in the data (dashed line; i.e., country-level intercepts).

**Figure S9.**
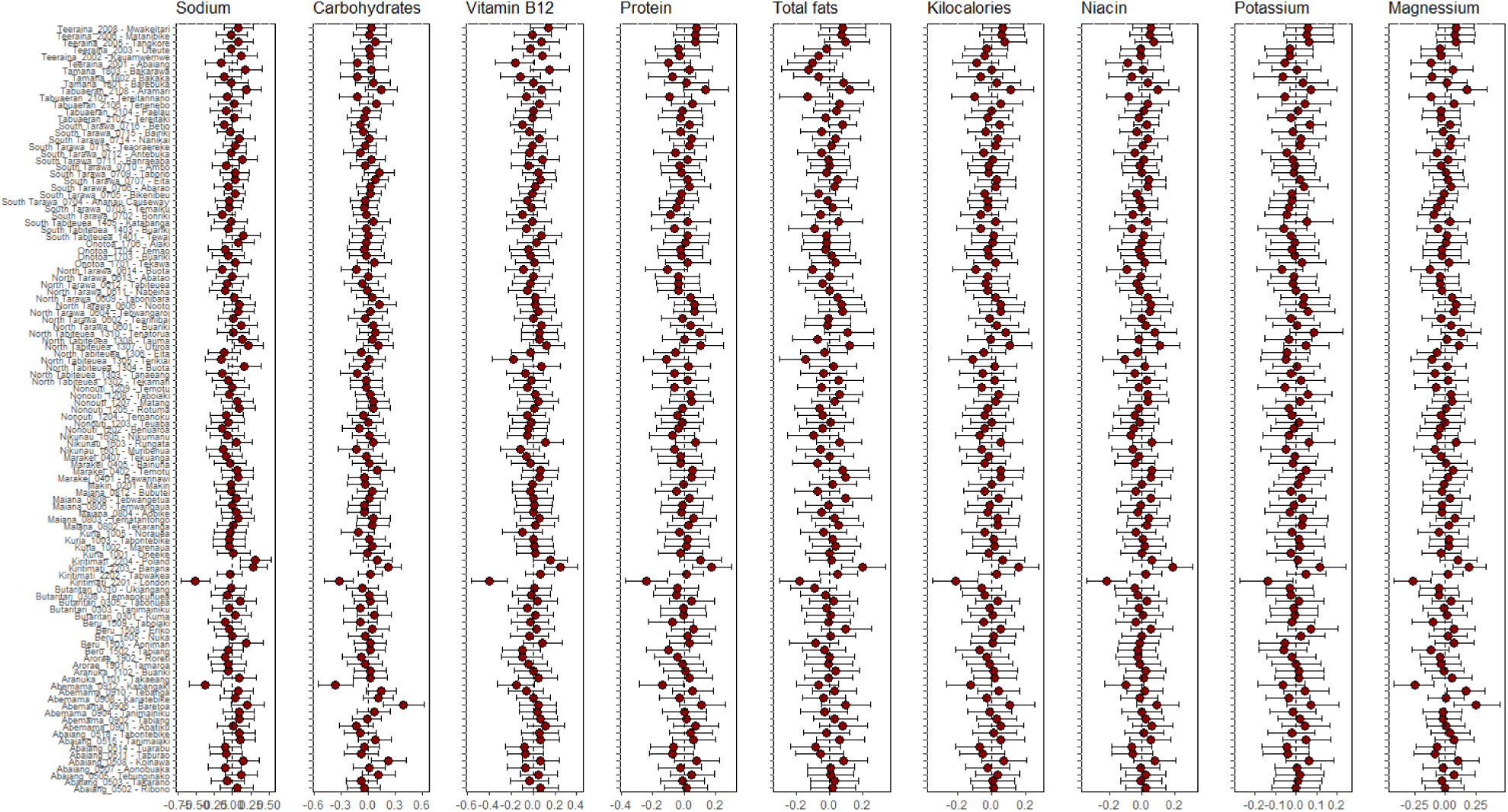
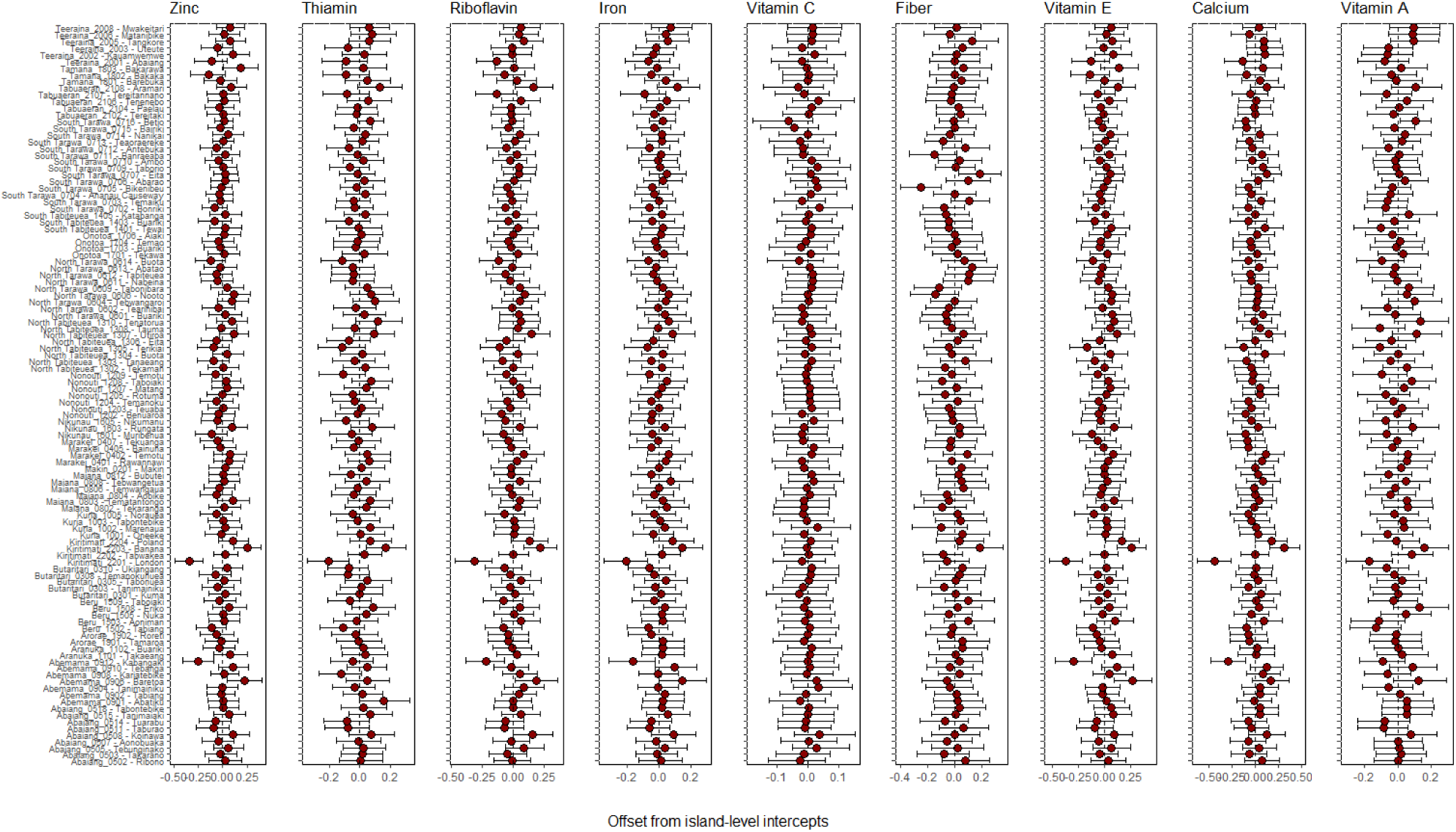
Community specific variability in nutrient intake adequacy met from aquatic foods as an offset from island-level estimated intercepts. Points are median estimates and intervals are 90% uncertainty intervals. Vertical dashed lines indicate island-level intercepts.

**Figure S10.**
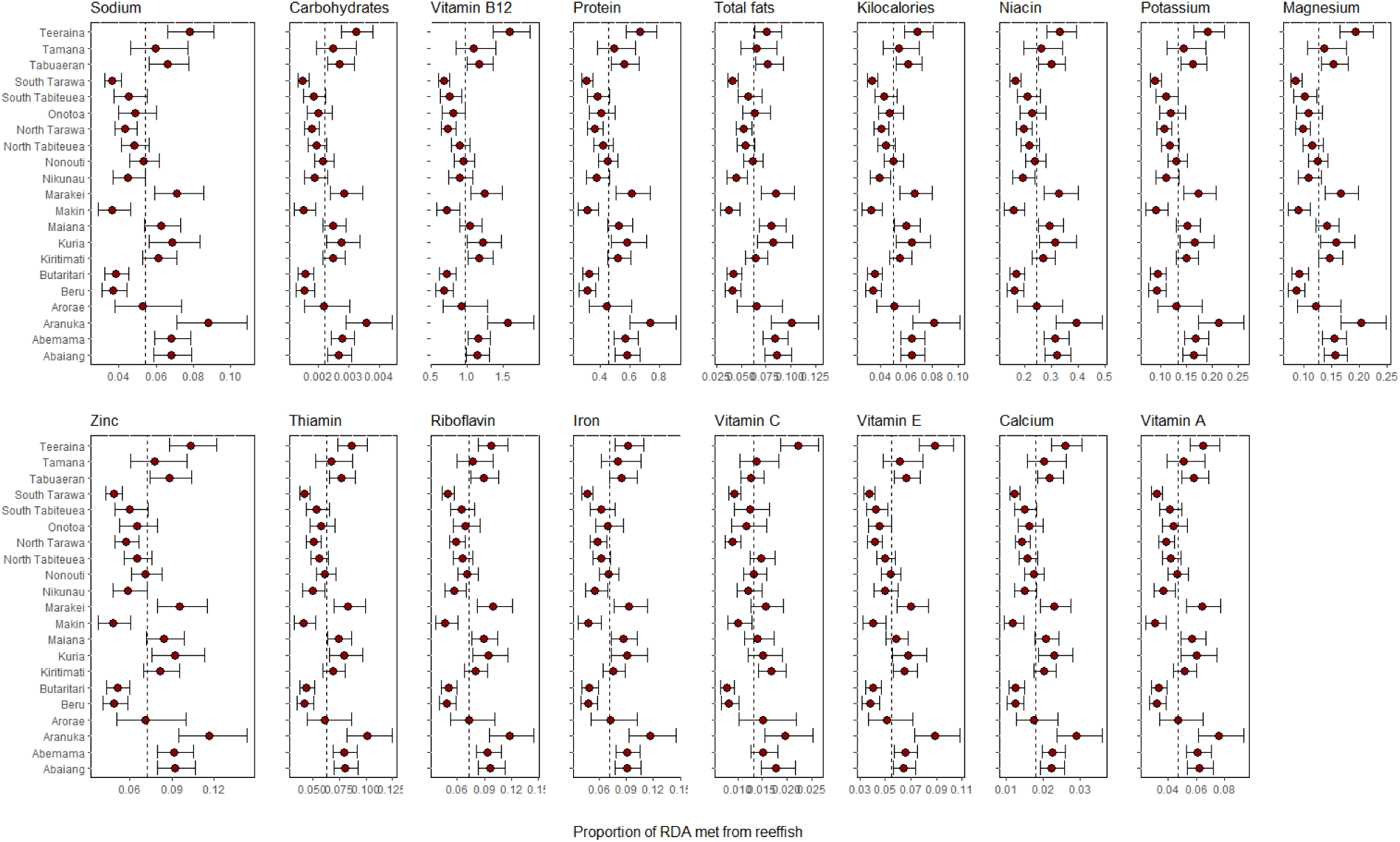
Island specific variability in nutrient intake adequacy from reef fish (proportion of nutrient-specific recommended daily allowance met from only reef fish). Points are median estimated nutritional adequacies and intervals are 90% uncertainty intervals. Vertical dashed line indicates average conditions for the entire population accounting for the nested structure in the data (dashed line; i.e., country-level intercepts). Note that the x axis varies across nutrients, and we did not include fibre because there was no variability (i.e., all values were zero).

**Figure S11.**
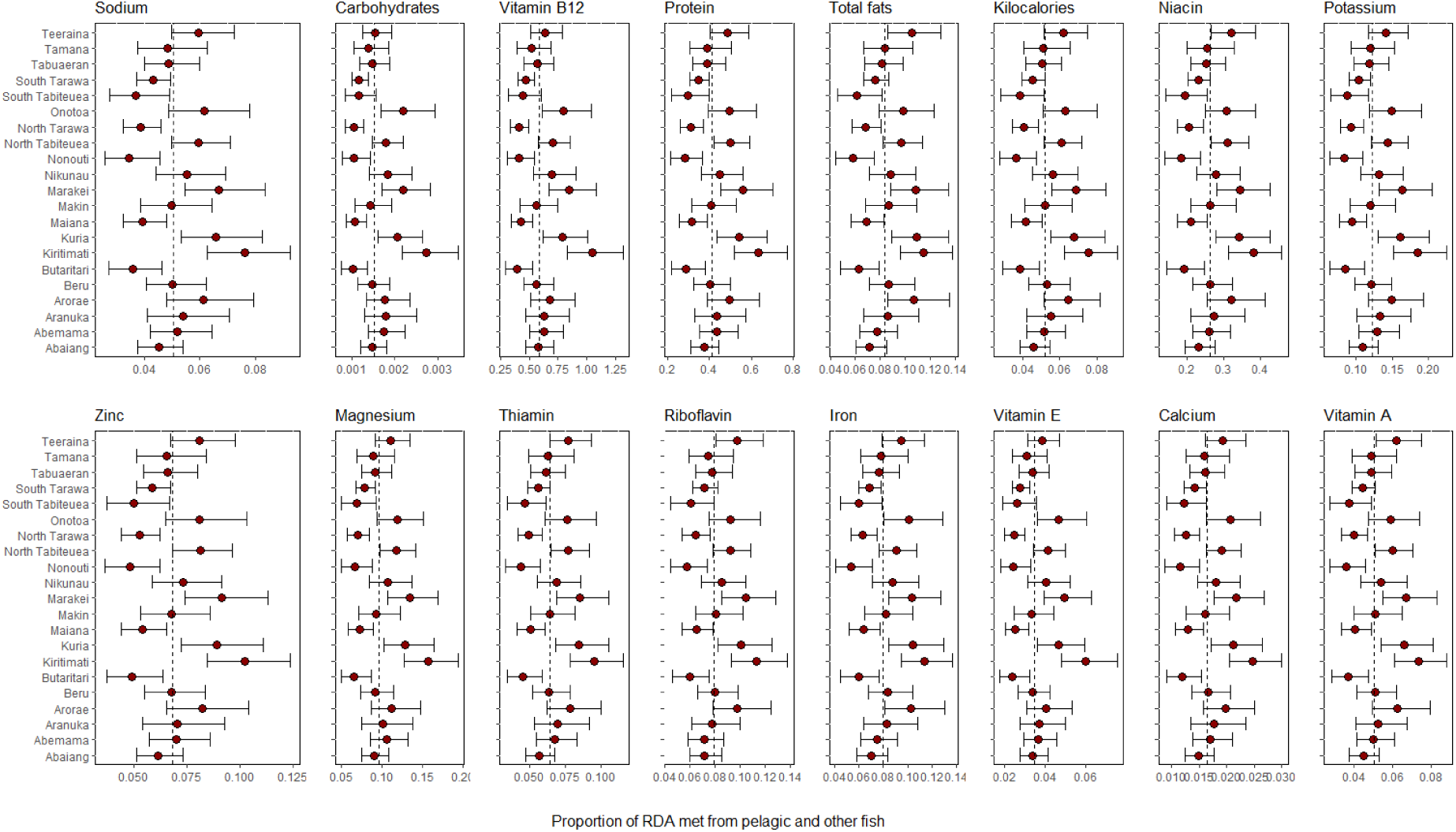
Island specific variability in nutrient intake adequacy from pelagic and other non-reef fish (proportion of nutrient-specific recommended daily allowance met from only pelagic and other fish). Points are median estimated nutrient intake adequacy and intervals are 90% uncertainty intervals. Vertical dashed line indicates average conditions for the entire population accounting for the nested structure in the data (dashed line; i.e., country-level intercepts). Note that the x axis varies across nutrients, and we did not include fibre or vitamin C because there was no variability (i.e., all values were zero).

**Figure S12.**
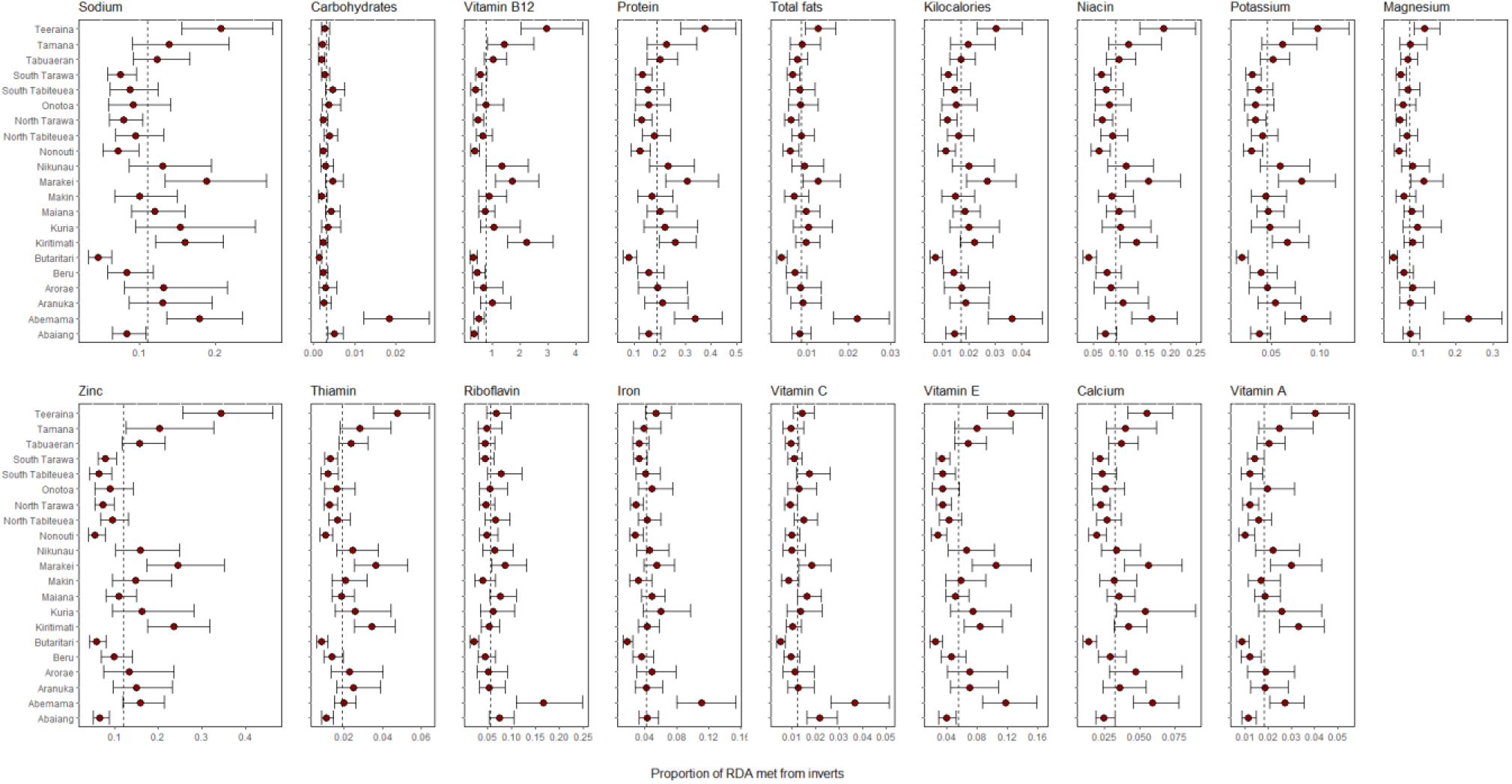
Island specific variability in nutrient intake adequacy from invertebrates (proportion of nutrient-specific recommended daily allowance met from only invertebrates). Points are median estimated nutrient intake adequacy and intervals are 90% uncertainty intervals. Vertical dashed line indicates average conditions for the entire population accounting for the nested structure in the data (dashed line; i.e., country-level intercepts). Note that the x axis varies across nutrients, and we did not include fibre because there was no variability (i.e., all values were zero).

**Figure S13.**
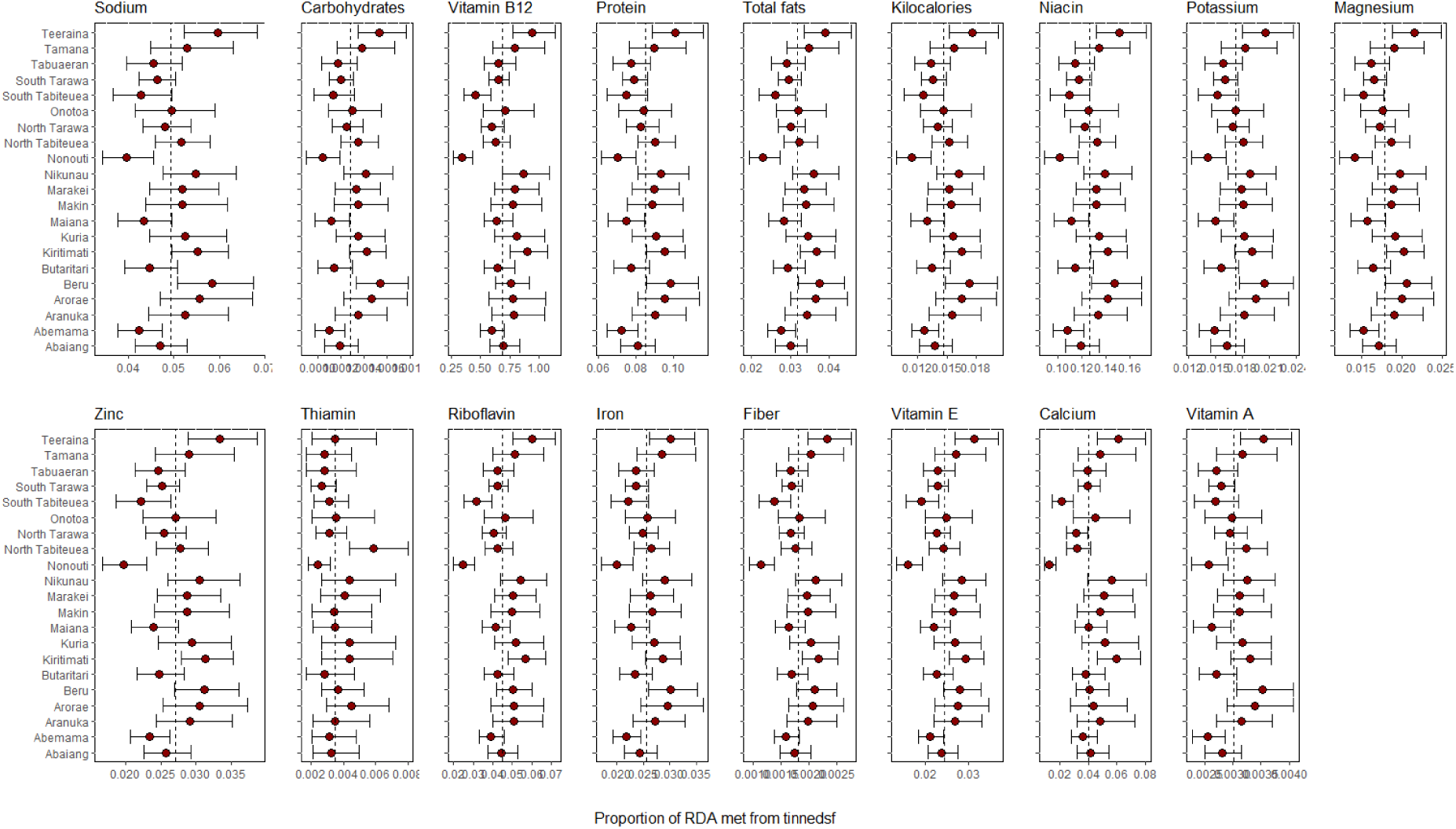
Island specific variability in nutrient intake adequacy from tinned seafood (proportion of nutrient-specific recommended daily allowance met from only tinned seafood). Points are median estimated nutrient intake adequacys and intervals are 90% uncertainty intervals. Vertical dashed line indicates average conditions for the entire population accounting for the nested structure in the data (dashed line; i.e., country-level intercepts). Note that the x axis varies across nutrients, and we did not include vitamin C because there was no variability (i.e., all values were zero).

**Figure S14.**
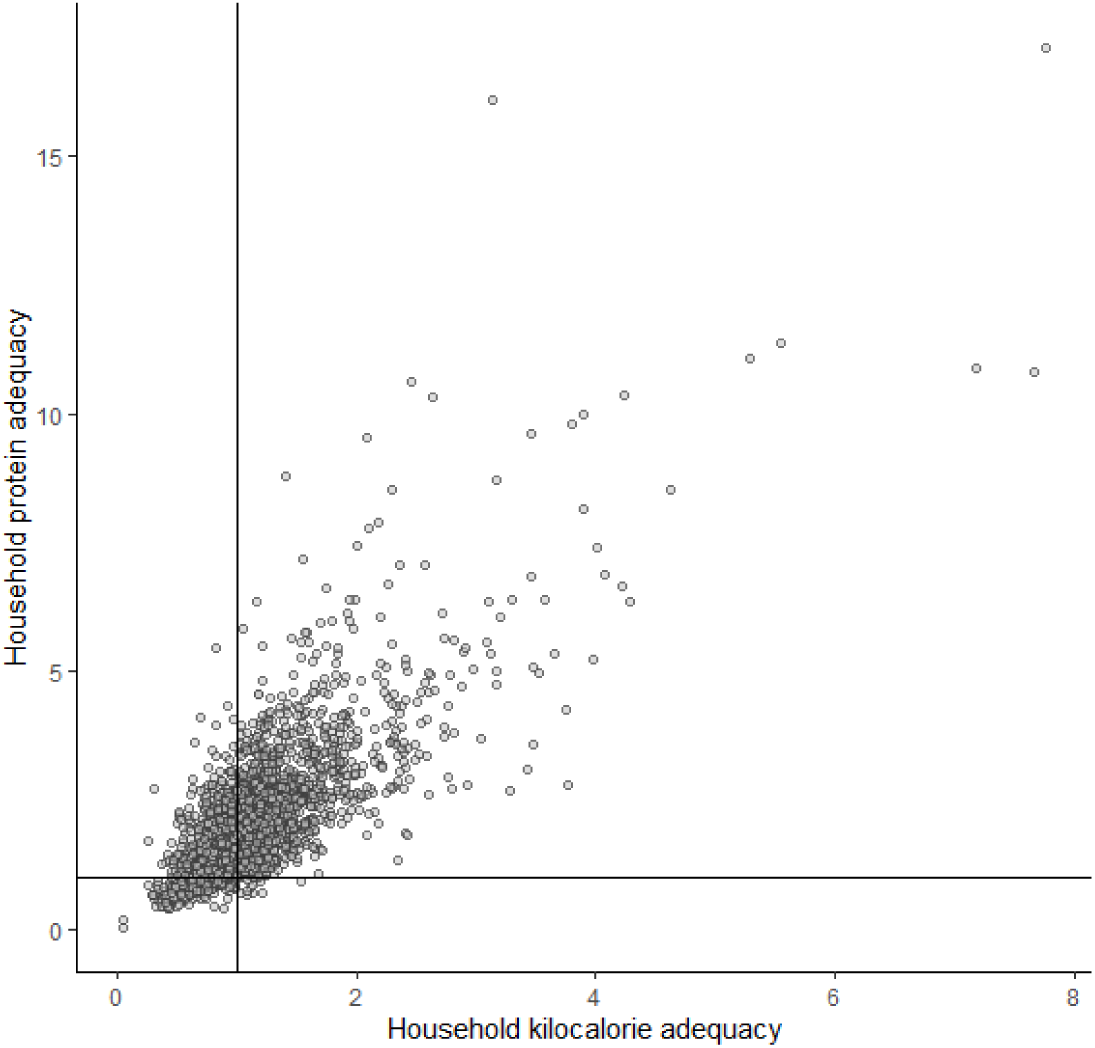
Relationship between household protein and kilocalorie adequacies. Each point is a household. Vertical and horizontal lines at one indicate the threshold of the household meeting or not kilocalorie and protein adequacies, respectively.

**Figure S15.**
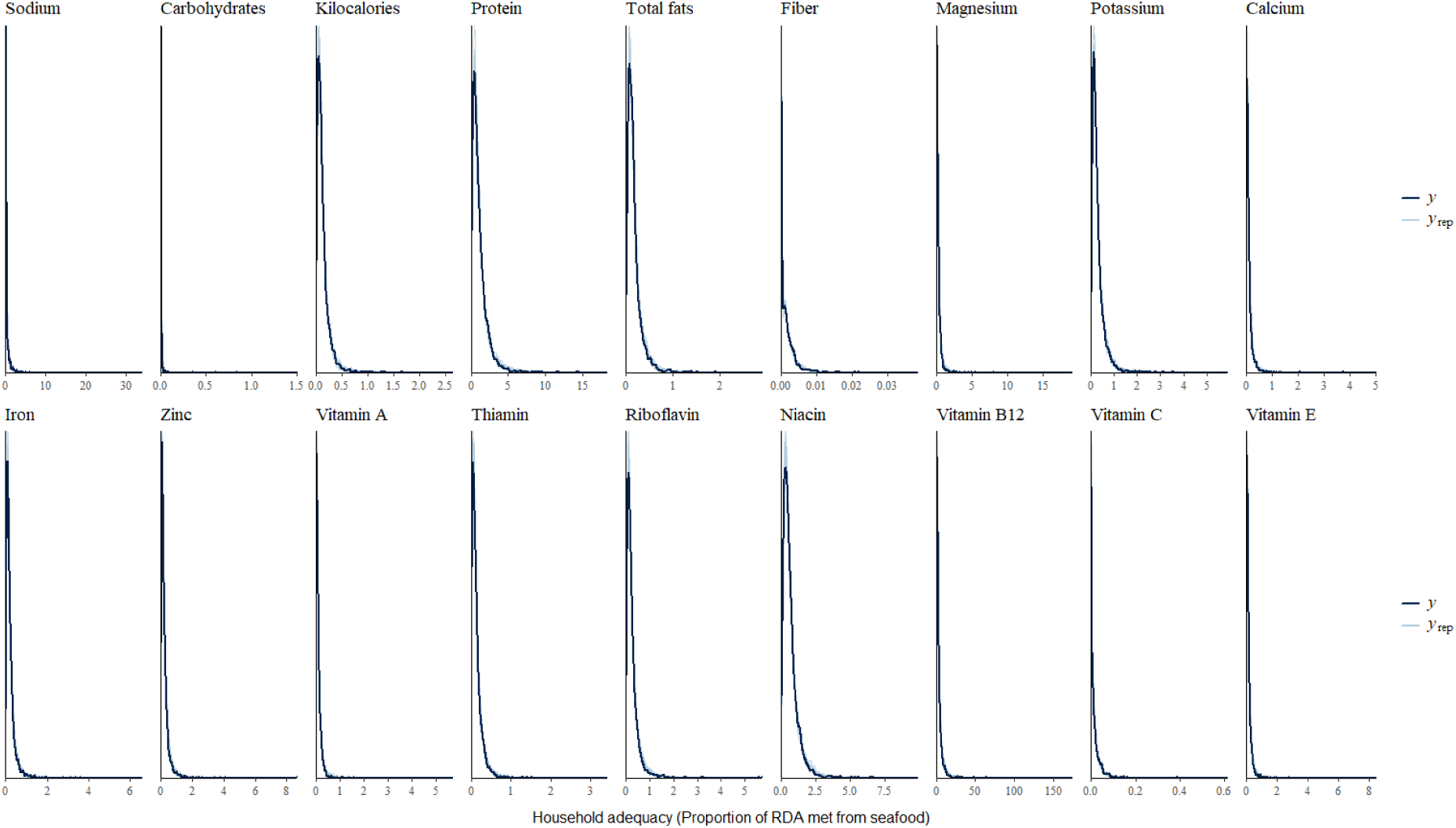
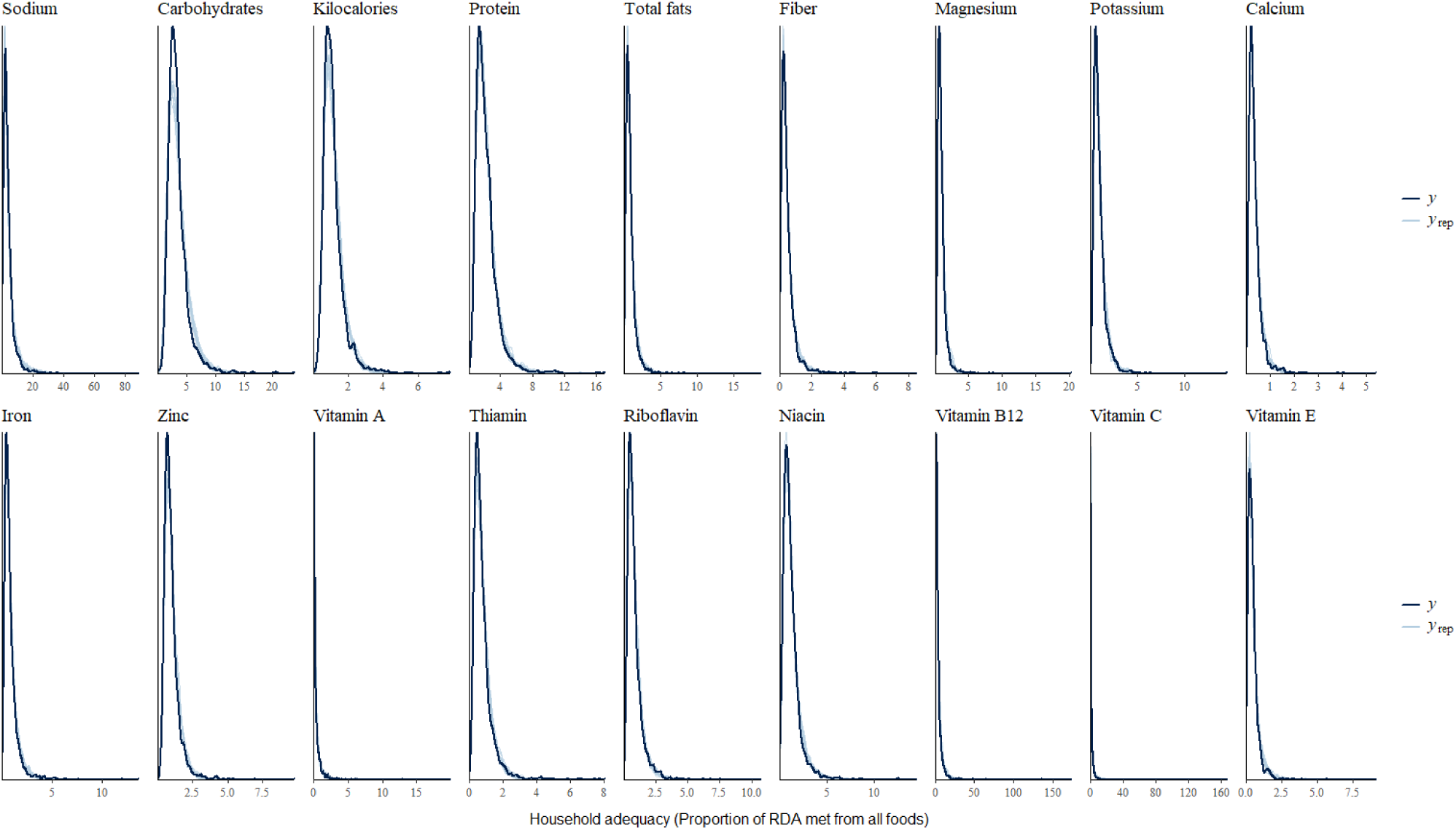
Example model fits. Black line is the observed distribution whereas light blue lines are posterior draws from the models.

## REFERENCES

1. Andrew, N. L., Allison, E. H., Brewer, T., Connell, J., Eriksson, H., Eurich, J. G., … & Tutuo, J. (2022). Continuity and change in the contemporary Pacific food system. Global Food Security, 32, 100608.

2. Atzori, D., Sonneveld, B. G., Alfarra, A., & Merbis, M. D. (2024). Nutrition fragility in isolation: Food insecurity in Small Island Developing States. Food Security, 16(2), 437–453.

3. Stevens, G. A., Beal, T., Mbuya, M. N., Luo, H., Neufeld, L. M., Addo, O. Y., … & Young, M. F. (2022). Micronutrient deficiencies among preschool-aged children and women of reproductive age worldwide: a pooled analysis of individual-level data from population-representative surveys. The Lancet Global Health, 10(11), e1590–e1599.

4. Passarelli, S., Free, C. M., Shepon, A., Beal, T., Batis, C., & Golden, C. D. (2024). Global estimation of dietary micronutrient inadequacies: a modelling analysis. The Lancet Global Health, 12(10), e1590–e1599.

5. Hawley, N. L., & McGarvey, S. T. (2015). Obesity and diabetes in Pacific Islanders: the current burden and the need for urgent action. Current diabetes reports, 15, 1–10.

6. Guell, C., Saint Ville, A., Anderson, S. G., Murphy, M. M., Iese, V., Kiran, S., … & Unwin, N. (2024). Small Island Developing States: addressing the intersecting challenges of non-communicable diseases, food insecurity, and climate change. The Lancet Diabetes & Endocrinology, 12(6), 422–432.

7. World Health Organization. (2023). Non communicable diseases. https://www.who.int/news-room/fact-sheets/detail/noncommunicable-diseases

8. Popkin, B. M., Adair, L. S., & Ng, S. W. (2012). Global nutrition transition and the pandemic of obesity in developing countries. Nutrition Reviews, 70(1), 3–21. 10.1111/j.1753-4887.2011.00456.x

9. FAO. 2023. The State of Food and Agriculture 2023 – Revealing the true cost of food to transform agrifood systems. Rome. 10.4060/cc7724en

10. Santos, J. A., McKenzie, B., Trieu, K., Farnbach, S., Johnson, C., Schultz, J., … & Webster, J. (2019). Contribution of fat, sugar and salt to diets in the Pacific Islands: A systematic review. Public health nutrition, 22(10), 1858–1871.

11. Ahern, M., Thilsted, S., Oenema, S., & Kühnhold, H. (2021). The role of aquatic foods in sustainable healthy diets. UN Nutrition Discussion Paper.

12. Farmery, A. K., Scott, J. M., Brewer, T. D., Eriksson, H., Steenbergen, D. J., Albert, J., … & Andrew, N. L. (2020). Aquatic foods and nutrition in the Pacific. Nutrients, 12(12), 3705.

13. Golden, C. D., Koehn, J. Z., Shepon, A., Passarelli, S., Free, C. M., Viana, D. F., … & Thilsted, S. H. (2021). Aquatic foods to nourish nations. Nature, 598(7880), 315–320.

14. Tigchelaar, M., Leape, J., Micheli, F., Allison, E. H., Basurto, X., Bennett, A., … & Wabnitz, C. C. (2022). The vital roles of blue foods in the global food system. Global Food Security, 33, 100637.

15. Tlusty, M. F., Tyedmers, P., Bailey, M., Ziegler, F., Henriksson, P. J., Béné, C., … & Jonell, M. (2019). Reframing the sustainable seafood narrative. Global Environmental Change, 59, 101991.

16. Hicks, C. C., Cohen, P. J., Graham, N. A., Nash, K. L., Allison, E. H., D’Lima, C., … & MacNeil, M. A. (2019). Harnessing global fisheries to tackle micronutrient deficiencies. Nature, 574(7776), 95–98.

17. Zamborain-Mason, J., Viana, D., Nicholas, K., Jackson, E. D., Koehn, J. Z., Passarelli, S., … & Golden, C. D. (2023). A decision framework for selecting critically important nutrients from aquatic foods. Current Environmental Health Reports, 10(2), 172–183.

18. Rimm, E. B., Appel, L. J., Chiuve, S. E., Djoussé, L., Engler, M. B., Kris-Etherton, P. M., … & Lichtenstein, A. H. (2018). Seafood long-chain n-3 polyunsaturated fatty acids and cardiovascular disease: a science advisory from the American Heart Association. Circulation, 138(1), e35–e47.

19. Seto, K. L., Friedman, W. R., Eurich, J. G., Gephart, J. A., Zamborain-Mason, J., Sharp, M., … & Golden, C. D. (2024). Characterizing pathways of seafood access in small island developing states. Proceedings of the National Academy of Sciences, 121(7), e2305424121.

20. O’Meara, L., Cohen, P. J., I’iluKafa, R., Wate, J. T., Albert, J., Bogard, J., … & Lam, F. V. (2023). Pacific food systems–The role of fish and other aquatic foods for nutrition and health. Apia, FAO.

21. United Nations. (2015). Transforming our world: the 2030 Agenda for Sustainable Development. Available at: https://sdgs.un.org/2030agenda

22. Golden, C. D., Gephart, J. A., Eurich, J. G., McCauley, D. J., Sharp, M. K., Andrew, N. L., & Seto, K. L. (2021). Social-ecological traps link food systems to nutritional outcomes. Global Food Security, 30, 100561.

23. FAO. 2020. The State of World Fisheries and Aquaculture 2020. Sustainability in action. Rome. 10.4060/ca9229en

24. Viana, D. F., Zamborain-Mason, J., Gaines, S. D., Schmidhuber, J., & Golden, C. D. (2023). Nutrient supply from marine small-scale fisheries. Scientific Reports, 13(1), 11357.

25. Bell, J. D., Kronen, M., Vunisea, A., Nash, W. J., Keeble, G., Demmke, A., Pontifex, S., & Andréfouët, S. (2009). Planning the use of fish for food security in the Pacific. Marine Policy, 33(1), 64–76. 10.1016/j.marpol.2008.04.002

26. Gombart AF, Pierre A, Maggini S. A Review of Micronutrients and the Immune System– Working in Harmony to Reduce the Risk of Infection. Nutrients. 2020;12(1):236. 10.3390/nu12010236

27. Gasperi, V., Sibilano, M., Savini, I., & Catani, M. V. (2019). Niacin in the central nervous system: an update of biological aspects and clinical applications. International journal of molecular sciences, 20(4), 974.

28. Abu-Ouf, N. M., & Jan, M. M. (2015). The impact of maternal iron deficiency and iron deficiency anemia on child’s health. Saudi medical journal, 36(2), 146.

29. Golden, C. D., Allison, E. H., Cheung, W. W., Dey, M. M., Halpern, B. S., McCauley, D. J., … & Myers, S. S. (2016). Nutrition: Fall in fish catch threatens human health. Nature, 534(7607), 317–320.

30. Bennett, A., Rice, E., Muhonda, P., Kaunda, E., Katengeza, S., Liverpool-Tasie, L. S. O., … & Gondwe, E. (2022). Spatial analysis of aquatic food access can inform nutrition-sensitive policy. Nature Food, 3(12), 1010–1013.

31. Scandurra, G., Romano, A. A., Ronghi, M., & Carfora, A. (2018). On the vulnerability of Small Island Developing States: A dynamic analysis. Ecological Indicators, 84, 382–392.

32. Hughes, T. P., Kerry, J. T., Baird, A. H., Connolly, S. R., Dietzel, A., Eakin, C. M., … & Torda, G. (2018). Global warming transforms coral reef assemblages. Nature, 556(7702), 492–496.

33. Pinsky M. L., Selden R. L., Kitchel Z. J. (2020). Climate-driven shifts in marine species ranges: scaling from organisms to communities. Ann. Rev. Mar. Sci. 12 (1), 153–179. doi: 10.1146/annurev-marine-010419-010916

34. Zamborain-Mason, J., Cinner, J. E., MacNeil, M. A., Graham, N. A., Hoey, A. S., Beger, M., … & Connolly, S. R. (2023). Sustainable reference points for multispecies coral reef fisheries. Nature Communications, 14(1), 5368.

35. Bell, J. D., Senina, I., Adams, T., Aumont, O., Calmettes, B., Clark, S., … & Williams, P. (2021). Pathways to sustaining tuna-dependent Pacific Island economies during climate change. Nature sustainability, 4(10), 900–910.

36. Eurich, J. G., Friedman, W. R., Kleisner, K. M., Zhao, L. Z., Free, C. M., Fletcher, M., … & Mills, K. E. (2024). Diverse pathways for climate resilience in marine fishery systems. Fish and Fisheries, 25(1), 38–59.

37. Popkin, B. M. (2017). Relationship between shifts in food system dynamics and acceleration of the global nutrition transition. Nutrition Reviews, 75(2), 73–82. 10.1093/nutrit/nuw064

38. Eme, P. E., Kim, N. D., Douwes, J., Burlingame, B., Foliaki, S., & Wham, C. (2020). Are households in Kiribati nutrition secure? A case study of South Tarawa and Butaritari. Food and nutrition bulletin, 41(1), 131–146.

39. Dietary Guidelines Advisory Committee. Scientific Report of the 2020 Dietary Guidelines Advisory Committee: Advisory Report to the Secretary of Agriculture and Secretary of Health and Human Services. U.S. Department of Agriculture, Agricultural Research Service; 2020. 10.52570/DGAC2020

40. Osendarp, S. J., Martinez, H., Garrett, G. S., Neufeld, L. M., De-Regil, L. M., Vossenaar, M., & Darnton-Hill, I. (2018). Large-scale food fortification and biofortification in low-and middle-income countries: a review of programs, trends, challenges, and evidence gaps. Food and nutrition bulletin, 39(2), 315–331.

41. Thow, A. M., Heywood, P., Schultz, J., Quested, C., Jan, S., & Colagiuri, S. (2011). Trade and the nutrition transition: strengthening policy for health in the Pacific. Ecology of food and nutrition, 50(1), 18–42.

42. Willett, W. (2012). Nutritional epidemiology. Oxford university press.

43. Troubat, N., & Sharp, M. K. (2021). Food consumption in Kiribati: Based on analysis of the 2019/20 Household Income and Expenditure Survey. Food & Agriculture Org.

44. Harris-Fry H, Shrestha N, Costello A, et al., Determinants of intra-household food allocation between adults in South Asia - a systematic review. Int J Equity Health. 2017;16:107. doi:10.1186/s12939-017-0603-

45. Eme, P. E., Burlingame, B., Douwes, J., Kim, N., & Foliaki, S. (2019). Quantitative estimates of dietary intake in households of South Tarawa, Kiribati. Asia Pacific journal of clinical nutrition, 28(1), 131–138.

46. Casey, E. M., Mojarrabi, M., Hannan-Jones, M. T., & Bogard, J. R. (2024). Measuring dietary intake in low-and middle-income countries: a systematic review of the methods and tools for estimating fish and seafood intake. Nutrition Reviews, 82(4), 453–466.

47. Campbell, B., & Hanich, Q. (2014). Fish for the future: Fisheries development and food security for Kiribati in an era of global climate change. WorldFish.

48. Golden, C. D., Ayroles, J., Eurich, J. G., Gephart, J. A., Seto, K. L., Sharp, M. K., … & Timeon, E. (2022). Study protocol: interactive dynamics of coral reef fisheries and the nutrition transition in Kiribati. Frontiers in Public Health, 10, 890381.

49. SPC; UOW; FAO. The Pacific Nutrient Database User Guide: A Tool to Facilitate the Analysis of Poverty, Nutrition and Food Security in the Pacific Region; Pacific Community, University of Wollongong and the Food and Agriculture Organization of the United Nations: Noumea, New Caledonia, 2020.

50. Statistics for Development Division Pacific Nutrient DataBase 2020 (PNDB 2020). Version Version 01 of the public-use dataset (July 2020) provided by the Pacific Data Hub - Microdata Library. Available online at: https://microdata.pacificdata.org/index.php/home

51. Institute of Medicine (US) Standing Committee on the Scientific Evaluation of Dietary Reference Intakes. Dietary Reference Intakes for Calcium, Phosphorous, Magnesium,Vitamin D, and Fluoride (1997); Dietary Reference Intakes for Thiamin, Riboflavin, Niacin, Vitamin B6, Folate, Vitamin B12, Pantothenic Acid, Biotin, and Choline (1998); Dietary Reference Intakes for Vitamin C, Vitamin E, Selenium, and Carotenoids (2000); Dietary Reference Intakes for Vitamin A, Vitamin K, Arsenic, Boron, Chromium, Copper, Iodine, Iron, Manganese, Molybdenum, Nickel, Silicon, Vanadium, and Zinc (2001); Dietary Reference Intakes for Energy, Carbohydrate, Fiber, Fat, Fatty Acids, Cholesterol,Protein, and Amino Acids (2002/2005); and Dietary Reference Intakes for Calcium and Vitamin D (2011). These reports may be accessed via www.nap.edu

52. Liu, A. G., Ford, N. A., Hu, F. B., Zelman, K. M., Mozaffarian, D., & Kris-Etherton, P. M. (2017). A healthy approach to dietary fats: understanding the science and taking action to reduce consumer confusion. Nutrition journal, 16, 1–15.

53. Kwon, Y. J., Lee, H. S., Park, J. Y., & Lee, J. W. (2020). Associating intake proportion of carbohydrate, fat, and protein with all-cause mortality in Korean adults. Nutrients, 12(10), 3208.

54. Stan Development Team, C. (2020). RStan: the R interface to Stan. R package version 2.21. 2.

55. Bürkner, P. C. (2017). brms: An R package for Bayesian multilevel models using Stan. Journal of statistical software, 80, 1–28.

56. Vehtari, A., Gelman, A., Simpson, D., Carpenter, B., & Bürkner, P. C. (2021). Rank-normalization, folding, and localization: An improved R for assessing convergence of MCMC (with discussion). Bayesian analysis, 16(2), 667–718.

57. Nesheim, M. C., Oria, M., Yih, P. T., & National Research Council. (2015). Dietary recommendations for fish consumption. In A framework for assessing effects of the food system. National Academies Press (US).

